# *In Vivo* Quantification of Histone Acetylation Turnover and Acetyl-CoA Sources Using ²H₂O Metabolic Labeling and High-Resolution Mass Spectrometry

**DOI:** 10.64898/2026.06.26.734905

**Authors:** Andrea Arias-Alvarado, Usman Sabir, Serguei Ilchenko, Spencer Parrish, Mirjavid Aghayev, Wentao He, Tsung-Heng Tsai, Guo-Fang Zhang, Takhar Kasumov

**Author notes:** Correspondence: Takhar Kasumov Department of Pharmaceutical Sciences, Northeast Ohio Medical University, Rootstown, OH 44272.

## Abstract

Dysregulated histone acetylation links cellular metabolism to gene expression, but measuring its *in vivo* turnover remains technically challenging. Here, we introduce a **^2^H2O**-based metabolic labeling method coupled with high-resolution Orbitrap mass spectrometry to quantify *in vivo* histone acetylation dynamics. The approach leverages differing deuterium incorporation rates between fast-labeling acetyl groups and slow-labeling peptide backbones. A two-tier analytical workflow uses full-scan mass spectrometry for mono-acetylated peptides, combined with parallel reaction monitoring (PRM) to resolve site-specific turnover and stoichiometry. Furthermore, monitoring acetyl-group plateau **^2^H** enrichment enables the evaluation of specific substrate contributions to the acetyl-CoA pool supporting histone acetylation.

To demonstrate biological utility, we applied this approach to mice maintained on a high-carbohydrate diet or subjected to 48-h fasting to assess nutrient-dependent histone acetylation dynamics. Acetyl-group labeling reflected the metabolic origin of acetyl-CoA, showing greater ^2^H enrichment in the fed state and reduced enrichment during fasting due to increased utilization of unlabeled fatty acid–derived acetyl-CoA. Fasting accelerated acetylation turnover across multiple histone sites and reduced overall acetylation stoichiometry. Quantitative tracing revealed that fatty acid oxidation becomes an important contributor to histone acetylation during fasting, whereas glucose remains the predominant source of nucleo-cytosolic acetyl-CoA (supplying > 60% of acetylation used carbon).

This approach enables simultaneous *in vivo* assessment of histone acetylation turnover, site occupancy, and acetyl-CoA substrate utilization, offering a robust platform to investigate metabolic-epigenetic crosstalk in health and disease.

## INTRODUCTION

The eukaryotic genome is organized into chromatin, a dynamic structure whose accessibility and function are governed by post-translational modifications (PTMs) of histone tails. Among these, lysine N(ε) acetylation (AcK) is a key regulator of transcriptionally active chromatin, promoting gene expression by neutralizing the positive charge of histones, weakening histone–DNA interactions, and facilitating recruitment of bromodomain-containing chromatin remodelers ^1–3^. In addition to functioning as an epigenetic mark, histone acetylation reflects cellular metabolic state, linking chromatin regulation to central metabolism ^4^. Acetylation is also widespread beyond histones, occurring on thousands of proteins including metabolic enzymes and transcription factors ^5–6^, highlighting its broad regulatory role. Consistent with this central role, dysregulated histone acetylation has been implicated in aging ^7^, dementia ^8^, cancer ^9^, metabolic diseases ^10^, and alcohol-associated liver disease ^11^.

Histone acetylation is highly dynamic, with stoichiometry governed by the balance between acetyl-CoA-dependent lysine acetyltransferases (KATs) and deacetylation by Zn²⁺-dependent histone deacetylases (class I, II, and IV HDACs) and NAD⁺-dependent sirtuins (class III HDACs). Transcriptionally active acetylation marks exhibit faster turnover than repressive modifications^12–13^, emphasizing the need for approaches that quantify acetylation dynamics rather than steady-state levels. However, most analytical methods measure acetylation occupancy rather than turnover, limiting insight into the dynamics of metabolic–epigenetic coupling *in vivo*. Moreover, site-specific measurements of acetylation turnover *in vivo* remain technically challenging, leaving the kinetic heterogeneity of histone acetylation across regulatory sites poorly defined.

Because acetylation depends directly on acetyl-CoA availability and cellular redox state, it functions as a metabolically responsive PTM linking nutrient status to gene regulation ^14^. Changes in energy substrate availability reshape acetyl-CoA pools and influence histone acetylation across tissues. For example, nutrient excess elevates nuclear–cytosolic acetyl-CoA and promotes histone acetylation and lipogenic gene expression ^15^. Conversely, although elevated β-hydroxybutyrate (BHB) acts as an endogenous HDAC inhibitor that can preserve histone acetylation in several tissues during fasting^16–17^, fasting and calorie restriction generally remodel the hepatic histone acetylation landscape and reduce overall hepatic histone acetylation ^18–19^. Under physiological conditions, glycolysis-derived pyruvate is a major source of acetyl-CoA for histone acetylation ^20^. When glucose availability is limited, alternative substrates, including fatty acids, acetate, and glutamine, can also contribute ^21^, highlighting the metabolic flexibility of acetyl-CoA production. Ketone bodies have generally been considered unable to support nucleo-cytosolic acetyl-CoA production for histone acetylation^22^; however, mitochondrial acetylcarnitine can be exported to the cytosol and regenerate acetyl-CoA, providing an alternative carbon source^23–24^. Consistent with this metabolic sensitivity, a high-fat diet reduces, while chronic alcohol exposure increases, acetylation of core histones in mouse liver ^10, 25–26^. Despite this metabolic flexibility, the relative contribution of these substrates to histone acetylation *in vivo* remains poorly defined.

Early measurements of acetylation turnover relied on radioactive acetate labeling, which lacked site-specific resolution ^27–28^. More recently, ^13^C tracers combined with mass spectrometry enabled site-specific acetylation measurements in cell culture ^29–34^, but these strategies have not been readily translated to *in vivo* systems. Other strategies, including SILAC-based proteomics ^35^ and label-free workflows ^32^, similarly remain restricted to *in vitro* studies. While recent stable-isotope approaches can relate PTMs to protein turnover ^36–37^, they do not directly quantify acetyl-group transfer rates or identify metabolic sources of acetyl-CoA used for acetylation.

Stable-isotope–coupled mass spectrometry studies indicate that histone acetylation is associated with increased histone turnover ^30^, consistent with acetylation-driven chromatin destabilization and enhanced nucleosome dynamics ^38^. Combinatorial acetylation of neighboring lysines further correlates with increased histone exchange, suggesting accelerated turnover of multi-acetylated H3 and H4 species. Although these findings highlight the interplay between acetylation and histone stability, they do not directly measure the turnover of the acetyl modification itself.

In this study, we developed a ²H₂O-based metabolic labeling strategy for *in vivo* quantification of histone acetylation turnover using high-resolution Orbitrap mass spectrometry. The method exploits differential ²H incorporation into the rapidly labeled acetyl group derived from acetyl-CoA versus the slowly labeled histone peptide backbone. Our analysis focuses on the canonical histones H3.1 and H4, targeting key regulatory lysine acetylation sites, H3.1K9, K14, K18, and K23, and H4K5, K8, K12, and K16, which recruit bromodomain-containing reader proteins39 to regulate chromatin accessibility, transcription, DNA repair, and autophagy, and are implicated in cancer and neurodegenerative diseases^39–42^. A two-tier analytical workflow integrates high-resolution full-scan mass spectrometry to quantify turnover of mono-acetylated peptides with parallel reaction monitoring (PRM) to resolve positional acetylation isomers and determine site-specific acetylation turnover and stoichiometry through diagnostic fragment ions. Measurement of plateau ²H enrichment in the acetyl moiety further enables estimation of the relative contributions of glycolytic and alternative substrates to the acetyl-CoA pool *in vivo*. Together, this strategy provides a quantitative framework to measure histone acetylation dynamics and link acetylation turnover to metabolic substrate utilization *in vivo*.

## EXPERIMENTAL SECTION

Detailed procedures are described in the Supporting Information.

### Materials

All materials used in this study are detailed in Supplementary Information.

#### Animal Experiments

All animal procedures were approved by the Northeast Ohio Medical University (NEOMED) Institutional Animal Care and Use Committee (IACUC) and conducted according to the Guidelines for the Care and Use of Laboratory Animals (National Institutes of Health publication 85-23). All animals were housed in a temperature-controlled facility under a 12-hour light/dark cycle with ad libitum access to standard chow and water.

#### Experimental Design

Male C57BL/6 wild-type mice (10 weeks old, 20–23 g; Jackson Laboratory) were used for all experiments. Method development was performed in chow-fed mice (n = 6). To assess substrate-dependent acetylation, an independent cohort (n = 12) was assigned to either chow or a high-carbohydrate diet (HCD; 70% carbohydrate, 35% sucrose) for three weeks **(Supplementary Table 1)**. During the final 48 h, chow-fed mice were fasted to promote fatty acid oxidation, whereas HCD-fed mice remained on diet, establishing metabolic states dominated by fatty acids or glucose, respectively.

#### ^2^H_2_O-metabolic Labeling

Mice were labeled with ²H₂O via an intraperitoneal bolus (28 µL/g body weight) followed by 8% ²H₂O in drinking water ^43^. Animals were euthanized at 0–24 h (0, 1, 2, 4, 8, and 24 h) to assess time-dependent incorporation of deuterium into histone acetyl groups.

#### Metabolic Phenotyping

During the final 24 h, mice were housed in metabolic cages (CLAMS) to assess food intake, activity, and respiratory exchange ratio. Body composition was measured by EchoMRI prior to euthanasia. Animals were anesthetized, blood was collected by cardiac puncture, and tissues were harvested, perfused with PBS, snap-frozen, and stored at −80 °C for subsequent analyses. Euthanasia was performed between 1–4 PM to minimize circadian variability ^44^.

##### Analytical procedures

###### Immunoblot Analysis

Liver lysates were prepared in RIPA buffer, and equal protein amounts (15 µg) were resolved by SDS-PAGE and transferred to PVDF membranes. After blocking, membranes were incubated with primary and HRP-conjugated secondary antibodies, and signals were detected and quantified using a FluorChem system. Antibody details are provided in the Supplementary Information.

###### Metabolic Profiling by GC-MS and LC-MS/MS

²H enrichment of body water ^45^, glucose ^46^, lactate ^47^, palmitate ^48^, acetylcarnitine ^49^, and acetyl-CoA ^50–51^ was quantified using established gas chromatography-mass spectrometry (GC–MS) and liquid chromatography tandem mass spectrometry (LC–MS/MS) methods with appropriate derivatization and internal standards. Detailed protocols and analytical parameters are provided in the Supplementary Information.

###### Histone Sample Preparation

Histones were acid-extracted and chemically derivatized with propionic anhydride to block unmodified lysines, followed by sequential trypsin/GluC digestion and peptide propionylation ^52^. Samples were purified and analyzed by LC–MS/MS. Detailed procedures are provided in the Supplementary Information.

###### High-Resolution Mass Spectrometry (DDA)

Proteomic analyses were performed using ultra-performance liquid chromatography (UHPLC) coupled to a Q Exactive Plus Orbitrap mass spectrometer operating in data-dependent acquisition (DDA) mode ^43^. Peptides were separated by nanoflow reverse-phase chromatography, followed by high-resolution MS1 scans and HCD-based MS/MS of the most abundant precursor ions. Detailed chromatographic and instrument parameters are provided in the Supplementary Information.

###### Protein/Peptide Identification

MS/MS spectra were searched using Mascot against the UniProt Mus musculus Reference Proteome (Proteome ID: UP000000589; release of June 10, 2026) database with a target–decoy strategy, applying a false discovery rate (FDR) ≤1% at peptide and protein levels. Enzyme specificity was set to trypsin and Glu-C, allowing a maximum of four missed cleavages. Relevant variable modifications, including propionylation and lysine acetylation and methylation (mono-, di-, and trimethylation) were included. Acetylated peptides were identified based on accurate mass and diagnostic fragment ions and were manually validated ^43^. Detailed search parameters and validation criteria are provided in the Supplementary Information. Annotated MS/MS spectra for all analyzed acetylated and methylated peptides are provided in **Supplementary Figures 1** and **2**.

**Figure 1.**
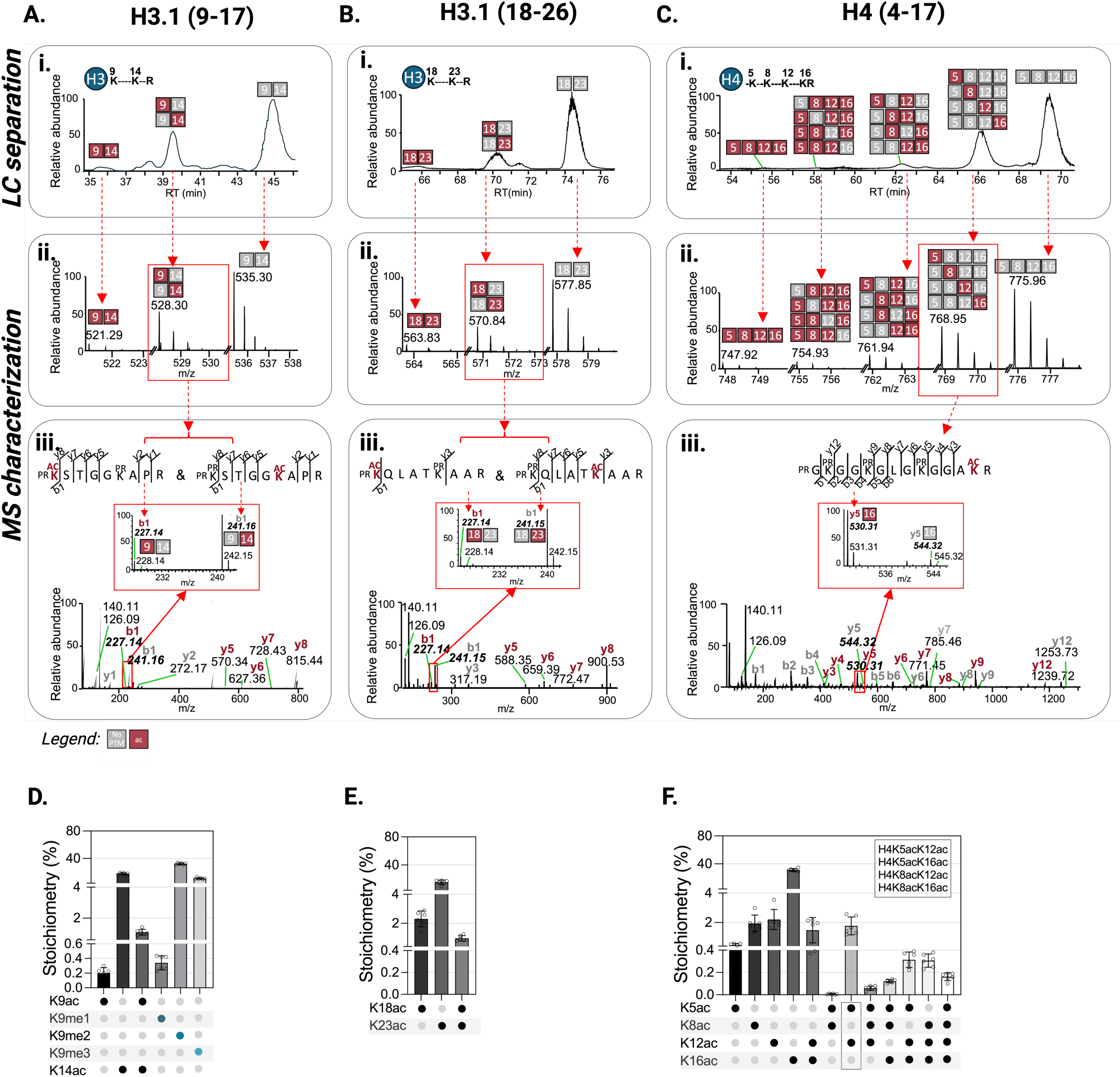
Two-tier mass spectrometry workflow for identification and quantification of histone acetylation. Histone H3.1 peptides ⁹KSTGGKAPR¹⁷ containing lysines K9 and K14 (9–17) (**A**) and ¹⁸KQLATKAAR²⁶ containing K18 and K23 (18–26) (**B**), and the histone H4 peptide ⁴GKGGKGLGGAKR¹⁷ containing K5, K8, K12, and K16 (4–17) (**C**), were analyzed after chemical derivatization and tryptic digestion. Prior to digestion, unmodified lysine residues were derivatized with propionic anhydride to generate propionylated lysines and prevent tryptic cleavage at internal lysine sites. This strategy generates peptides with identical amino acid sequences but distinct acetylation states, enabling quantitative analysis of native, mono-, di-, tri-, and tetra-acetylated species by high-resolution mass spectrometry. (i) Representative chromatographic separation of positional acetylation isomers in histone H3 and H4 peptides: H3.1 (9–17 and 18–26) and H4 (4–17). Chemical derivatization with propionic anhydride enables separation of native and fully acetylated peptide species, whereas mono-, di-, and tri-acetylated positional isomers co-elute chromatographically (**A–C**), requiring MS/MS fragment-ion analysis for site-specific discrimination. Gray and red squares denote unmodified and acetylated sites, respectively. (ii) **MS1-based quantification of global acetylation states.** Full-scan MS1 spectra capture composite signals from co-eluting positional isomers, resulting in average ^2^H-isotopic labeling kinetics that reflect ensemble acetylation turnover. Although MS1-level measurements distinguish between native and fully acetylated peptide species based on mass differences, they lack the resolution to differentiate positional acetylation isomers. (iii) **Site-specific acetylation analysis using targeted parallel reaction monitoring (PRM).** Diagnostic ***b*** and ***y*** fragment ions enable discrimination of positional acetylation isomers and accurate quantification of acetylation at individual lysine residues. (**D-F**) Acetylation stoichiometry of mono- and di-acetylated H3.1 peptides (**D**: 9–17 and **E**: 18–26), and di-, tri-, and tetra-acetylated H4 peptides (**F**: 4–17), determined from fragment-ion intensities. In panels **A–B**, the insets show magnified ***b1*** fragment ions corresponding to endogenously acetylated and chemically propionylated forms, which were used to determine the relative abundances of H3K9ac, H3K14ac, H3K18ac, and H3K23ac. In panel **Ciii**, the inset displays a magnified ***y5*** fragment ions (indicated by an arrow) corresponding to K16ac along with its non-acetylated fragment ions, which were used to quantify site-specific acetylation. The H3.1 peptide spanning residues 9–17 also contains mono-, di-, and tri-methylation at lysine 9 (see **Supplementary** Figure 1).

**Figure 2.**
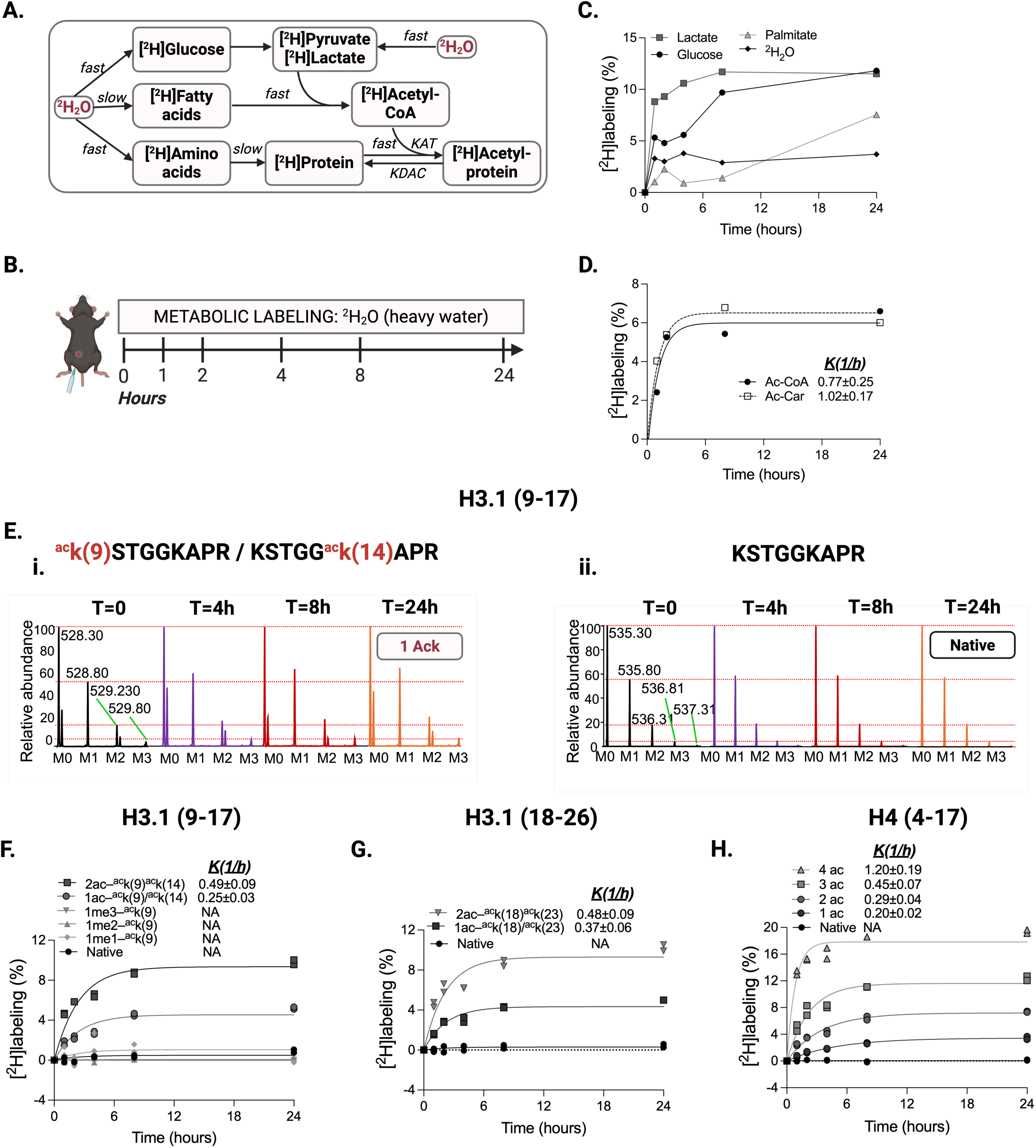
MS1-based ²H-metabolic labeling for quantifying histone acetylation turnover and acetyl-CoA sources *in vivo*. **(A) Principle of the ²H₂O labeling strategy.** Rapid exchange between body water and metabolic intermediates leads to incorporation of deuterium into acetyl-CoA, primarily via glycolytic substrates (glucose → pyruvate). Because histone acetylation turnover occurs within minutes to hours whereas histone protein turnover occurs over days to weeks, short-term (≤24 h) labeling selectively reports ²H incorporation into acetyl groups rather than peptide backbones. Slower labeling of fatty acid–derived acetyl-CoA enables inference of acetyl-CoA metabolic sources. **(B) Experimental design.** Mice received an intraperitoneal bolus of ²H₂O followed by continuous access to ²H₂O-enriched drinking water for up to 24 h. Animals were euthanized at defined time points to quantify time-dependent incorporation of deuterium into histone acetyl groups. **(C)** Time-dependent ²H enrichment of body water and selected metabolites (glucose, lactate, and palmitate) measured by mass spectrometry. **(D)** ²H labeling kinetics of acetyl-CoA and acetyl-carnitine measured by LC–MS/MS. **(E)** Representative high-resolution MS1 spectra of isobaric mono-acetylated H3.1 (9–17) peptides (K9ac or K14ac) (Figure 2Ei) and the corresponding native peptide (Figure 2Eii). Progressive enrichment of M1–M3 isotopomers in acetylated peptides reflects ²H incorporation into the acetyl moiety, whereas the native peptide shows no isotopic shift. (**F–H**) Time-dependent ²H incorporation into acetylated histone peptides quantified from MS1 isotopomer distributions of mono- and di-acetylated H3.1 (9–17) (**F**), mono- and di-acetylated H3.1 (18–26) (**G**), and mono-, di-, tri-, and tetra-acetylated H4 (4–17) peptides (**H**). Labeling increases proportionally with acetyl group number, enabling estimation of apparent acetylation turnover rates. Native (**F-H**) and methylated (**F**) peptides showed no detectable ²H enrichment.

###### Parallel Reaction Monitoring (PRM) Analysis

Site-specific acetylation turnover was quantified by PRM on a Q Exactive Plus, providing enhanced sensitivity and quantitative accuracy for fragment-ion–based isotope analysis ^53^. A multiplexed method with optimized HCD fragmentation (stepped NCE 25–45%) and narrow precursor isolation (3 m/z quadrupole isolation window with a 0.4 m/z offset) was used to minimize co-isolation interference^54^. Target peptides included histone H3.1 ⁹KSTGGKAPR¹⁷ and ¹⁸KQLATKAAR²⁶, and histone H4 ⁴GKGGKGLGKGGAKR¹⁷, monitored using inclusion lists defined by precursor m/z (±10 ppm), charge state, and retention-time windows derived from prior DDA runs (**Supplementary Table 2**). Detailed acquisition parameters are provided in the Supplementary Information.

###### Calculation of Site-specific Histone Acetylation Stoichiometry

Following propionylation, all lysines were fully acylated, and positional isomers with identical acetyl/propionyl content co-eluted with indistinguishable MS1 masses, precluding MS1-level resolution (**Figure 1Ai-ii, Bi-ii, and Ci–Cii**). Site-specific acetylation was therefore determined using diagnostic *b* and *y* ions in MS/MS spectra. Histone PTMs were quantified with EpiProfile 2.0 ^55^, with additional peptides incorporated based on Mascot-identified retention times, and all assignments were manually validated in QualBrowser (Thermo Fisher Scientific, version 3.0.63).

Stoichiometry was calculated by pairing endogenous acetylated fragment ions with their propionylated counterparts and expressing each isomer as a fraction of total signal, as described by Feller *et al.*^56^; the same approach was used for H3K9 methylation. Site-specific abundances for monoacetylated H3.1 (⁹KSTGGKAPR¹⁷; ¹⁸KQLATKAAR²⁶) and H4 (⁴GKGGKGLGKGGAKR¹⁷) peptides were derived from diagnostic MS2 ions and scaled to MS1 precursor signals. Di- and triacetylated H4 species were quantified using site-specific y ions, while the fully acetylated peptide was confirmed by multiple diagnostic fragments. Certain diacetylated isomers requiring MS3 ^56^ were not resolved and are reported as combined values. The quadruply acetylated H4K5acK8acK12acK16ac peptide was chromatographically resolved and confirmed by multiple site-diagnostic *b* and *y* ions (**Supplementary Figure 2Bv, Supplementary Table 3**). Detailed calculations are provided in the Supplementary Information.

###### Mass Isotopomer Distribution Analysis

Peptides and fragment-ion isotopologues were quantified manually using Xcalibur Qual Browser (Thermo Fisher Scientific). Extracted ion chromatograms were generated for acetylated and native histone H3 and H4 peptides. Isotopic distributions were determined with mass tolerances of ±10 ppm for full-scan MS1 and ±0.1 m/z for MS2 fragment ions. Because ²H₂O-based turnover measurements are highly sensitive to spectral accuracy, kinetic analyses were restricted to acetylated peptides with high signal intensity (>10⁵ arbitrary units), thereby minimizing uncertainty associated with low-abundance ions. Only modification sites meeting this criterion were included in turnover calculations.

Incorporation of ²H into the acetyl moiety redistributes the isotopic envelope, resulting in a progressive decrease in the relative abundance of the monoisotopic peak (M0) and a corresponding increase in heavier isotopomers (M1 and M2). In full-scan data, the fractional abundance of the monoisotopic isotopomer at time *t*, E₀(t), was calculated as:

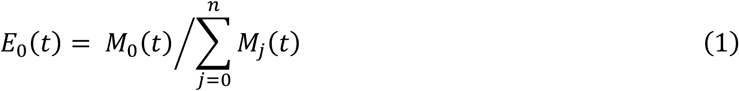

For fragment-ion–specific analysis, isotopomer abundances were quantified by integrating the m/z values of the corresponding *b*- and *y*-ions. Although PRM increased scan depth (40–200 scans), fragment-ion isotopic envelopes were often truncated, with reliable detection limited to M0 and M1; therefore, ²H enrichment was calculated using only these two isotopomers.

Because incorporation of ²H shifts signal from the monoisotopic peak to heavier isotopomers, total ^2^H labeling at time *t*, E(t), was expressed as:

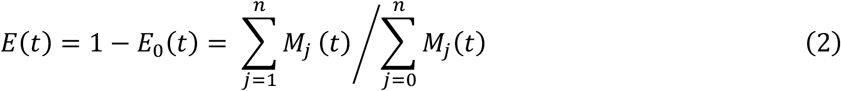

When fragment ions could not selectively resolve site-specific acetylation, site-specific ^2^H enrichment was inferred from differences in total labeling between fragment ions. For example, the *y7* ion of H4 (4–17) of di- and triacetylated species contain contributions from both K12 and K16; therefore, ^2^H enrichment at K12ac was calculated by subtracting the labeling of *y5* (representing K16) from that of *y7* (**Supplementary Figure 2Biii-iv**).

###### Acetylation Turnover Analysis

Our acetylation kinetic calculations assume a metabolic steady state in which acetylation and deacetylation rates are equal. The turnover rate constant (*k*) was estimated using a one-compartment model by fitting an exponential growth function to the time-dependent ²H labeling, *E(*t), of intact tryptic peptides or PRM fragment ions after subtraction of the natural background enrichment:

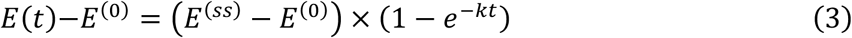

where *E*^(0)^ and *E*^(*ss*)^ are the baseline (pre-labeling) and steady-state (plateau) enrichments, respectively.

The acetylation half-life (*t1/2*) was calculated based on the turnover rate constant *k* as:

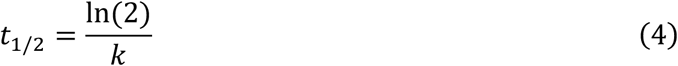

###### Statistical Analysis

Time-dependent incorporation of deuterium from ²H₂O into histone-bound acetyl groups was analyzed using established precursor–product kinetic models. Fractional synthesis rates (FSR), acetylation rate constants (*k*), and turnover half-lives (*t*₁*/*₂) were estimated by nonlinear least-squares fitting of labeling curves to mono-exponential or appropriate compartmental models. Fasting-induced changes in acetylation turnover were evaluated in a site-specific manner by jointly modeling time-course data from all acetylated peptides containing the same lysine residue. The parameters *k*_HCD_ and *k*_Fast_ represent site-level turnover rate constants under high-carbohydrate diet (HCD) and fasting (Fast) conditions, respectively. Turnover rate estimates are reported as mean ± SEM. Differences in turnover rates were assessed using *F*-tests comparing a model with a shared turnover rate constant (k) and condition-specific ^2^H enrichments to a model with condition-specific turnover rate constants and plateau enrichments. Differences in plateau ^2^H enrichment were similarly evaluated by comparing a model with condition-specific turnover rate constants but a shared plateau enrichment to a model with both condition-specific turnover rate constants and plateau enrichments. In the differential analysis, changes in turnover rates were considered statistically significant at *P* < 0.05, with significant changes indicating altered acetylation kinetics between dietary states. Model performance was evaluated by residual analysis and R². All statistical analyses, graph generation, and kinetic curve fitting were performed using GraphPad Prism (version 11.0.0; GraphPad Software, San Diego, CA, USA).

## RESULTS AND DISCUSSION

### Site-specific Histone Acetylation Stoichiometry in Mouse Liver

Histone tails contain multiple closely spaced lysine residues that undergo extensive acetylation and methylation. Site-specific acetylation analysis is complicated by multiple neighboring lysine residues that generate chromatographically unresolved, isobaric acetylated peptides. While ion mobility mass spectrometry provides partial separation of positional isomers, most key mono- and diacetylated H4 isomers could not be resolved in full-scan spectra ^57^. In contrast, fragment-ion–based prognostic ions analysis has enabled resolution of combinatorial acetylation sites^56^. In bottom-up histone proteomics, trypsin cleaves unmodified but not modified lysine and arginine residues, necessitating chemical derivatization of unmodified lysines to generate uniform tryptic peptides ^58^. Although ¹³C- or ²H-labeled acetic anhydrides generate chemically equivalent derivatives for stoichiometric analysis ^59^, their mass shifts (2.007 and 3.019 Da) overlap with M2 and M3 isotopomers. This overlap confounds stable isotope tracer–based measurements, particularly ²H metabolic labeling, which alters isotopomer distributions without producing discrete mass shifts ^60–62^. A doubly labeled ^13^C₄D₆-acetic anhydride has been used for cell-culture studies ^31, 33^, but its high cost and generation of complex, chromatographically overlapping acetylated species limit sequence coverage and quantitative accuracy. Therefore, we derivatized unmodified lysine residues with propionic anhydride ^63^. Although propionylated and acetylated peptides may exhibit modest differences in ionization efficiency, their chemical distinction enables partial chromatographic separation of mono-, di-, tri-, and tetra-acetylated species (**Figure 1Ai, 1Bi, and 1Ci**), reducing analytical complexity and improving both site-specific stoichiometric and tracer-based quantification.

We identified 33 histone proteoforms (**Supplementary Table 4**), including core histones (H2A, H2B, H3, and H4), the linker histone H1, and multiple variants, confirmed by unique peptides, present in both acetylated and methylated forms in wild type mouse liver. For instance, H2AX was acetylated at K5 and K36, whereas H2A type 3 showed acetylation at K127 and K129. A single acetylation site (K116) was detected in H2B type 1-C/E/G, while H2B type 3-B was methylated at K12. Additional acetylation and methylation sites were observed across linker histone isoforms H1.0, H1.1, H1.3, H1.4, and H1.5. Acetylation was most extensive in histones H3.1, a replication-dependent canonical H3 variant, and H4, with enrichment at H4K16ac, a key regulator of higher-order chromatin structure ^38^. H3.1 was acetylated at K9, K14, K18, K23, K36, K64, and K79, with concurrent mono-, di-, or trimethylation at K9, K27, and K36. In addition to the highly abundant K16 acetylation, histone H4 displayed acetylation at K5, K8, K12, and K79, while K20 was methylated. (**Supplementary Table 4**).

Given their central roles in metabolic gene regulation and their enrichment in our datasets, subsequent analyses were focused on histones H3.1 and H4. To quantify acetylation dynamics, we analyzed H3.1 peptides ^9^KSTGGKAPR^17^, ^18^KQLATKAAR^26^, and the H4 peptide ^4^GKGGKGLGKGGAKR^17^, each containing multiple acetylation sites and collectively generating 6 H3.1 and 15 H4 acetylated species. Propionylation of unmodified lysines and N-terminal amines enabled chromatographic separation of peptides by acetylation state but not by positional isomers (**Figure 1Ai, 1Bi, and 1Ci**). In contrast, mono-, di-, and tri-methylated H3K9 forms were partially resolved chromatographically and distinguished from native and acetylated forms based on MS1 (**Supplementary Figure 1A**) and MS2 spectra (**Supplementary Figure 1B-F**). Although positional mono-acetylated isomers of H3.1 peptides ^9^KSTGGKAPR^17^ (K9ac vs K14ac) and ^18^KQLATKAAR^26^ (K18ac vs K23ac) co-eluted and were indistinguishable at the MS1 level (**Figure 1Aii** and **1Bii**), site-diagnostic MS2 fragment ions enabled unambiguous resolution and site-specific quantification (**Figure 1Aiii** and **1Biii**). Specifically, H3K9ac and H3K14ac were resolved using *b1* and *y8* ions (**Figure 1Aiii**), and the same MS2-based strategy distinguished H3K18ac and H3K23ac in ^18^KQLATKAAR^26^ using *b1* and *y7* ions, respectively (**Figure 1Biii**). For H4, mono- , tri-, and adjacent di-acetylated isomers (K5acK8ac and K12acK16ac) were unambiguously characterized based on MS2 spectra (**Supplementary Figure 2**) and quantified using *y5*, *y7*, and *y12* ions (**Figure 1Ciii** and **Supplementary Figure 2Bii-iv**, see supplementary methods). In contrast, non-adjacent di-acetylated species (K5acK12ac, K5acK16ac, K8acK12ac, and K8acK16ac) were reported as combined abundances.

Quantitative PTM stoichiometry analysis encompassing native, acetylated, and methylated species revealed that in wild-type mouse liver under a normal chow diet, H3K14 displayed the highest acetylation stoichiometry (17.9 ± 1.3%), whereas H3K9 was predominantly methylated (32.3 ± 1.3% and 11.3 ± 1.4% for di- and trimethylation, respectively.) (**Figure 1D**). Substantial acetylation was also observed at H3K23 (15.4%), while H3K9 and H3K18 acetylation were minimal (∼0.2% and ∼2%, respectively) (**Figure 1D-E**). Both diacetylated H3 peptides exhibited low stoichiometry, with ∼1% for K9K14 and <1% for K18K23. In H4, K16 showed the highest acetylation stoichiometry (31.3±1.9%), followed by K12, K8, and K5, with multiple low-abundance di- and tri-acetylated species and a trace (∼0.2%) quadruply acetylated form (**Figure 1F**). Together, these results provide site-resolved acetylation stoichiometry of histones H3.1 and H4, and methylation stoichiometry of histone H3.1, in mouse liver, offering a quantitative basal reference for subsequent kinetic and stable-isotope tracer–based analyses of histone acetylation dynamics under physiological and metabolic perturbations.

### Rationale for ²H₂O-Based Histone Acetylation Turnover Measurements

We used ²H₂O metabolic labeling approach to quantify acetylation turnover by measuring ²H incorporation into the acetyl moiety of acetylated peptides (**Figure 2A**). Administered in drinking water, ²H₂O serves as a safe, non-radioactive, and inexpensive universal tracer to quantify lipid, protein, DNA, and RNA synthesis in free-living organisms, including humans^64–68^. Following administration, ^2^H2O rapidly equilibrates with total body water, including intracellular compartments, providing a uniform precursor pool. Deuterium incorporation depends on the number of metabolically exchangeable carbon-bound hydrogens and the turnover rate of the labeled molecule ^69^. Recently, we demonstrated that ^2^H2O labeling can detect acetylation-dependent differences in protein stability and turnover^25, 43^.

The present approach extends ^2^H2O metabolic labeling to the quantification of histone acetylation turnover. This application relies on several well-established assumptions. First, body-water enrichment rapidly reaches isotopic steady state and serves as a stable source of deuterium for the synthesis of both amino acids and acetyl-CoA^70^. Previous studies have validated rapid labeling and steady-state enrichment of tissue amino acid acetyl-CoA precursor pools during ^2^H2O administration^68, 70^. Second, deuterium incorporated into acetyl-CoA is transferred quantitatively to downstream acetylated products. Direct measurements have demonstrated incorporation of body-water-derived deuterium into hepatic acetyl-CoA and its subsequent transfer to newly synthesized lipids and other biosynthetic products^43^. Importantly, the methyl hydrogens of the acetyl moiety are carbon-bound and do not undergo significant hydrogen–deuterium exchange under physiological conditions^71^. Consequently, deuterium incorporated into acetyl-CoA is retained during enzymatic transfer of the acetyl group to lysine residues, supporting the use of acetyl-group labeling to assess acetylation turnover.

Under ²H₂O exposure, the isotopic enrichment of an acetylated peptide reflects two processes: incorporation into peptide backbones via protein synthesis and incorporation into acetyl groups via ²H-labeled acetyl-CoA (**Figure 2A**). Our analytical strategy exploits the marked difference in turnover rates between these processes. Histone acetylation turnover occurs on the scale of minutes to hours ^30, 52^, whereas histone protein turnover spans weeks ^62, 72^. Therefore, short-term (≤24 h) ²H₂O labeling preferentially captures labeling of the acetyl moiety while minimizing contributions from peptide backbone synthesis. Combined with high-resolution Orbitrap mass and fragment-ion analysis, this approach enables direct quantification of site-specific *in vivo* acetylation turnover in H3.1 (residues 9–17 and 18–26) and H4 (residues 4–17) peptides while simultaneously resolving positional acetylation isomers as described above.

### MS1-Based Analysis of Histone Acetylation Turnover and Acetyl-CoA Sources Using ²H₂O

To establish a ²H₂O-based approach for simultaneous quantification of histone acetylation turnover and acetyl-CoA substrate contributions *in vivo*, wild-type mice on a chow diet were labeled for up to 24 h using an initial ²H₂O bolus (28 µL/g body weight), followed by 8% ²H₂O in drinking water (**Figure 2B**), achieving ∼3.5% steady-state body water enrichment (**Figure 2C**). To assess metabolic sources of acetyl-CoA based on established precursor–product relationships^73^, we quantified time-dependent ²H labeling of plasma glucose, lactate (a stable surrogate for glycolytic pyruvate that is in rapid equilibrium with pyruvate and can be analyzed alongside palmitate by single-step TBDMS derivatization), palmitate, acetyl-CoA, acetyl-carnitine, and the acetyl group of K14 in the H3.1 (9-17) peptide (KSTGGKAPR) **(Figure 2C-E)**.

During short-term (≤24 h) ²H₂O tracer exposure, acetyl-group ²H enrichment primarily reflects glycolytic substrates, as glucose and pyruvate turn over substantially faster than fatty acids^74^. Consistently, glucose and lactate labeled rapidly, with lactate reaching isotopic plateau within ∼8 h, whereas palmitate labeled slowly and exhibited substantially lower enrichment (**Figure 2C**). The higher lactate enrichment relative to glucose and palmitate likely reflects rapid glycolytic turnover and isotopic equilibration, whereas glucose enrichment is buffered by continuous dietary input and endogenous production, and palmitate enrichment remains low due to its slow turnover.

To assess the feasibility of using ²H₂O labeling to quantify acetyl-CoA source and turnover, we measured the ^2^H-labeling of acetyl-CoA and its proxy metabolite acetyl-carnitine, which is in rapid equilibrium with acetyl-CoA and has recently been shown to serve as a source of acetyl groups for histone acetylation^23^. Both metabolites labeled rapidly, with turnover rate constants of 0.77±0.25 h⁻¹ and 1.02±0.17 h⁻¹, respectively, and reached nearly identical isotopic plateau enrichments within 4 h of labeling (**Figure 2D**). Their enrichment was lower than that of glycolytic intermediates (glucose and lactate) but substantially higher and appeared earlier than palmitate labeling, indicating that the acetyl-CoA pool turns over rapidly and is predominantly supplied by glycolytic carbon through mitochondrial pyruvate oxidation rather than by fatty acid oxidation.

We next quantified ²H incorporation into native and mono-acetylated H3.1 (9-17) peptides. Because K9ac and K14ac positional isomers were unresolved by chromatography or MS1 (**Figure 1A**), they were analyzed as a combined population. Full-scan MS1 spectra showed progressive enrichment of M1 and M2 isotopomers in acetylated peptides (**Figure 2Ei)**, whereas native peptides remained unlabeled (**Figure 2Eii)**, indicating selective ²H incorporation into acetyl groups rather than histone backbones (**Figure 2E-F**). Because precursor acetyl-CoA labeling reached a stable enrichment very rapidly, then we used ^2^H-labeling of acetylated histone to quantify acetylation turnover. The labeling kinetics of acetylated histones were well fit by a first-order exponential model with no systematic deviations. While we cannot exclude minor fluctuations in acetylation flux, these observations support the use of a quasi-steady-state approximation for estimating turnover rate constants^73^. Consistently, H3.1 peptides methylated at K9 site showed no detectable ²H incorporation (**Figure 2F**), reflecting the lack of hydrogen exchange from S-adenosylmethionine–derived methyl groups and slow protein turnover. Acetyl-histone ²H enrichment (**Figure 2F**) followed the expected precursor–product relationship, being lower than that of upstream metabolites (glucose, lactate, and acetyl-CoA) but substantially higher than palmitate (**Figure 2C–D**). This pattern indicates that histone acetyl groups are preferentially derived from glycolytic substrates rather than fatty acids, consistent with prior ¹³C-tracer studies^52^. While the present approach provides a quantitative assessment of glycolytic versus non-glycolytic substrate utilization, future integration with substrate-specific ^13^C tracers may enable more detailed mapping of individual acetyl-CoA sources.

Supporting the validity of the ^2^H₂O-labeling approach, the di-acetylated H3.1 (9-17) peptide (K9acK14ac) exhibited approximately twofold higher plateau enrichment (∼20%) than mono-acetylated species (∼10%), as expected for two independently labeled acetyl groups (**Figure 2F**). Kinetic modeling yielded apparent turnover rate constants of 0.25 ± 0.03 h⁻¹ for mono-acetylated and 0.49 ± 0.09 h⁻¹ for di-acetylated peptides, corresponding to acetylation half-lives of ∼1–2 h, orders of magnitude faster than histone protein turnover^10^. Although direct comparisons with *in vivo* mouse data are not available, these estimates are consistent with the rapid acetylation dynamics reported in cell-based studies^30, 33, 52^ and support a higher overall acetylation flux in diacetylated species^30, 33^.

Similarly, rapid ^2^H-labeling was observed for H3.1 ¹⁸KQLATKAAR²⁶ peptides, with mono- and di-acetylated species showing progressive and approximately doubled labeling, yielding turnover rates of 0.37 ± 0.06 and 0.48 ± 0.09 h⁻¹, respectively (**Figure 2G, Supplementary Figure 3A**). The accelerated turnover of diacetylated peptides compared to mono-acetylated species may arise from their lower abundance (**Figure 1D-E**), requiring increased turnover rates to sustain functional occupancy at these sites. Because MS1 analysis does not resolve acetylation sites within unresolved mono-acetylated peptides, these values represent composite rather than site-specific rates.

As discussed above, acetylation of neighboring lysine residues 5, 8, 12, and 16 within the histone H4 peptide ⁴GKGGKGLGKGGAKR¹⁷ generates multiple mono-, di-, tri-, and tetra-acetylated species, with all but the tetra-acetylated form comprising unresolved positional isomers, thereby complicating site-specific acetylation turnover analysis. Like histone H3.1, no detectable ²H enrichment was observed in the native peptide, whereas mono-acetylated peptide mixtures showed a gradual, time-dependent increase in ²H enrichment in MS1 spectra (**Supplementary Figure 3B**). As anticipated, the magnitude of ²H enrichment increased proportionally with the number of acetyl groups, from one to four (**Figure 2H)**. Exponential fitting revealed correspondingly higher apparent rate constants, reflecting the cumulative contribution of multiple acetylation sites. These findings are consistent with previous reports linking greater acetylation occupancy to increased acetylation flux^30, 33^ (**Table 1**).

**Table 1.**
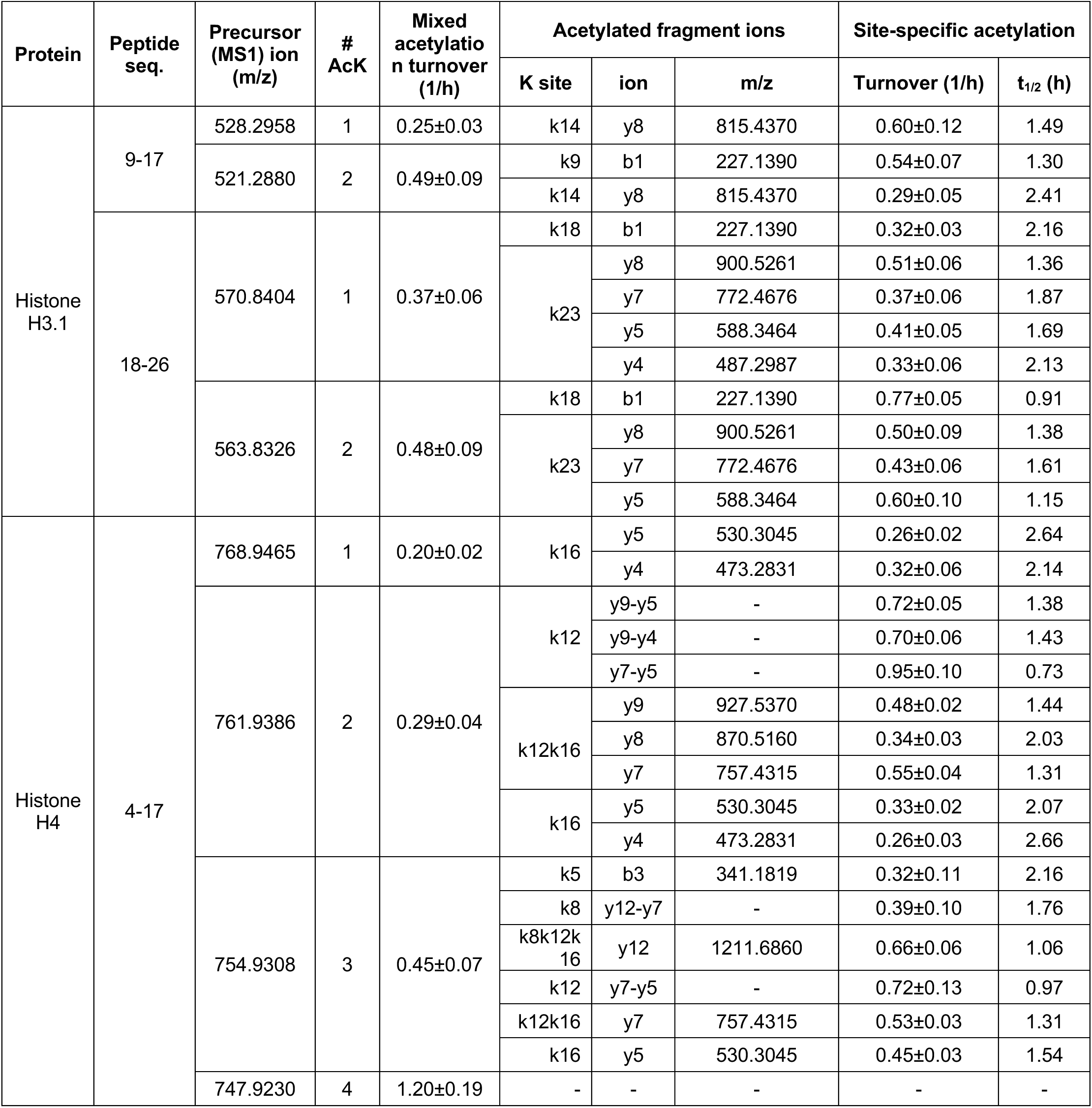
Quantified acetylated histone peptides and diagnostic fragment ions used for site-specific acetylation turnover analysis by a two-tier mass spectrometry workflow. The table lists acetylated histone H3.1 and H4 peptides quantified at the MS1 level together with diagnostic MS/MS fragment ions used for positional assignment of acetylated lysine residues. Acetylation turnover was determined using a two-tier strategy combining MS1 isotopomer analysis of acetylated peptides with fragment-ion–resolved MS/MS measurements, enabling resolution of positional acetylation isomers and quantification of site-specific acetylation turnover during ²H₂O metabolic labeling.

Collectively, these data indicate that over short labeling duration (≤24 h), ²H incorporation into histone peptides predominantly reflects acetyl-group turnover rather than protein synthesis. The close correspondence between acetyl-group labeling and glucose/lactate kinetics supports glycolytic substrates as the dominant source of acetyl-CoA driving histone acetylation *in vivo*, while highlighting the limitation of MS1-based analysis for resolving site-specific acetylation kinetics.

### Site-Specific Acetylation Turnover by Fragment-Ion–Resolved ²H₂O Labeling

#### H3.1(9-17) and (18-26) peptides

To measure site-specific acetylation turnover, we exploited the high resolving power of the Orbitrap to quantify diagnostic fragment ions from the mixed mono-acetylated H3.1 (18-26) peptides containing acetylation at either K18 or K23. PRM analysis distinguished the two isomers based on site-specific *b* and *y* ions. Specifically, the *y4, y5, y7*, and *y8* ions uniquely report H3K23ac, whereas the *b1* ion uniquely reports H3K18ac (**Supplementary Figure 1Biii**). As an illustration of this approach, high-resolution MS2 spectra of the diagnostic *y7* and *b1* ions showed progressive enrichment of M1 and M2 isotopomers, reflecting ²H incorporation into the acetyl groups at H3K23ac and H3K18ac, respectively (**Figure 3A**). Because these ions contain only a single acetylated lysine residue, their isotopomer distributions provide a direct, site-resolved measure of acetylation turnover. One-phase exponential fitting of labeling kinetics yielded turnover rate constants of 0.37 ± 0.06 h⁻¹ for H3K23ac (*y7*) and 0.32 ± 0.03 h⁻¹ for H3K18ac (*b1*), corresponding to apparent half-lives of 1.9 and 2.2 h, respectively (**Figure 3B**; **Table 1**). Although no additional *b* ions were available to independently validate H3K18ac turnover, analysis of the *y4, y5*, and *y8* fragment ions yielded a consistent H3K23ac half-life of 1.7±0.3 h, supporting the robustness of the site-specific turnover measurements (**Supplementary Figure 4A; Table 1**). In addition, *y5* and *y8* fragment-ions analysis enabled site-specific measurement of acetylation turnover in the diacetylated H3.1 (18–26) peptide, revealing faster apparent turnover than the corresponding monoacetylated species (**Figure 3C; Supplementary Figure 4B).**

**Figure 3.**
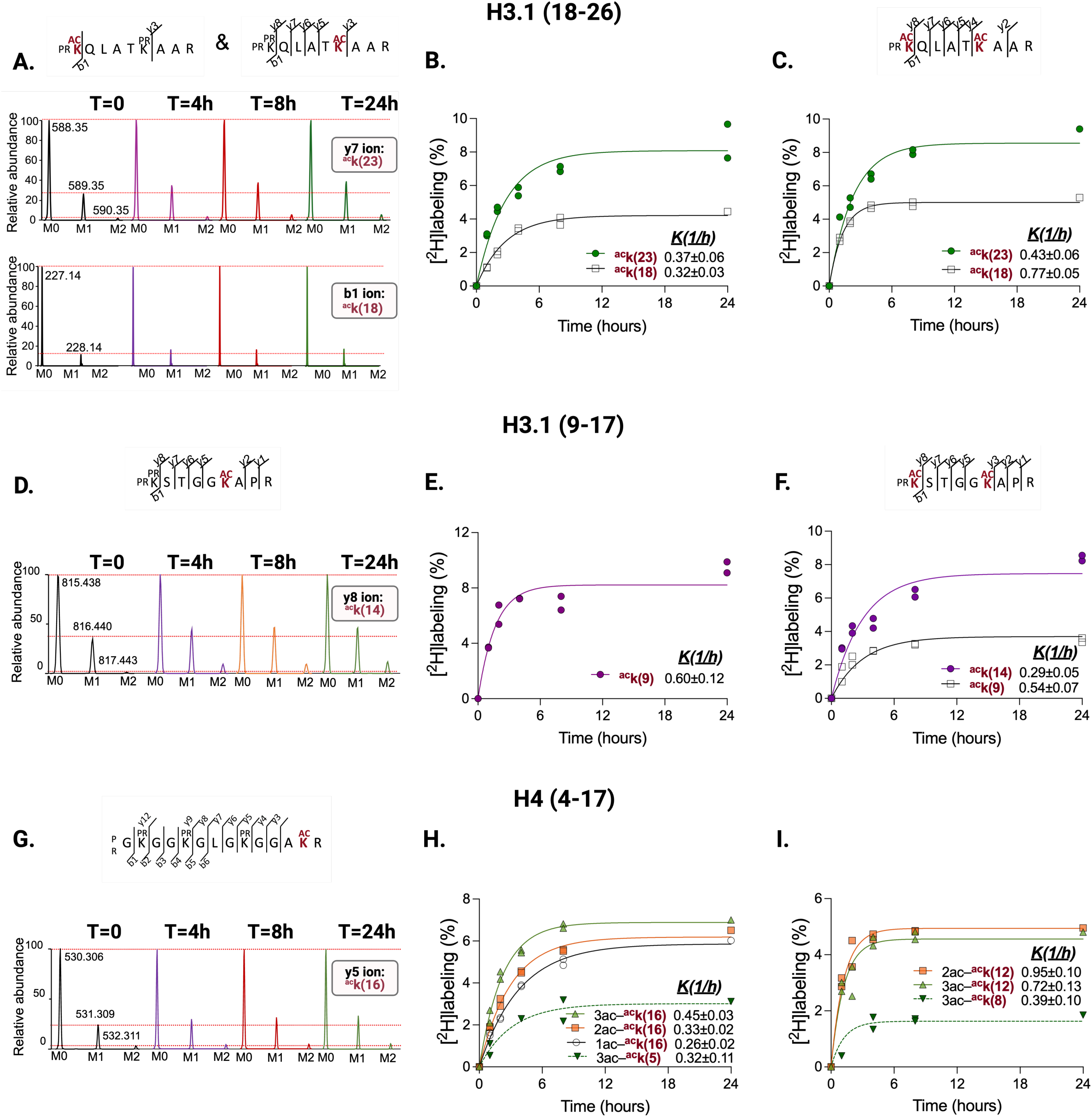
MS2 (fragment-ion)–based ²H-metabolic labeling for quantification of site-specific histone acetylation turnover *in vivo*. **(A)** High-resolution MS2 spectra of diagnostic fragment ions from mono-acetylated H3.1 (18–26) peptides acetylated at K23 (top) and K18 (bottom). Progressive enrichment of M1–M2 isotopomers in fragment ions containing the acetyl group reflects ²H incorporation into the acetyl moiety at the corresponding lysine residues. (**B–C**) Time-dependent ²H incorporation into diagnostic fragment ions enables site-specific acetylation turnover measurements. Labeling of fragment ions containing the acetyl group was quantified for mono- and di-acetylated H3.1 (18–26) peptides (**B–C**). (**D**) High-resolution MS2 spectra of diagnostic fragment ions from mono-acetylated H3.1 (9–17) peptides acetylated at K14. Progressive enrichment of M1–M2 isotopomers in fragment ions containing the acetyl group reflects ²H incorporation into the acetyl moiety at the corresponding lysine residues. (**E–F**) Time-dependent ²H incorporation into diagnostic fragment ions enables site-specific acetylation turnover measurements. Labeling of fragment ions containing the acetyl group was quantified for mono- and di-acetylated H3.1 (9–17) peptides. **(G)** High-resolution MS2 spectra of diagnostic fragment ions from mono-acetylated H4 (4–17) peptides acetylated at K16. Progressive enrichment of M1–M2 isotopomers in fragment ions containing the acetyl group reflects ²H incorporation into the acetyl moiety at the corresponding lysine residues. **(H)** Time-dependent ²H incorporation into diagnostic ***y5*** fragment ions from mono-, di-, and tri-acetylated H4 (4–17) peptides and ***b3*** fragment ions from the tri-acetylated species enables quantification of acetylation turnover at H4K16 and H4K5. Increased labeling with higher acetylation states reflects enhanced acetylation turnover in polyacetylated H4 peptides. **(I)** Estimation of acetylation turnover at H4K8 and H4K12 derived from calculated ²H enrichment differences between positional fragment ions containing these residues.

A similar fragment-ion–resolved approach enabled site-specific quantification of K14 acetylation turnover from the mono-acetylated H3.1 (9–17) peptide, yielding a turnover rate of 0.60±0.12 h⁻¹ (**Figure 3D-E**). Although low *b1* signal intensity precluded reliable estimation of low stoichiometry (∼0.2%) K9ac turnover, comparison with mixed MS1 kinetics and site-resolved K14ac measurements suggest that K9ac turnover occurs on a similar timescale, consistent with dynamic acetylation at transcriptionally active H3 residues. In addition, fragment-ion analysis further allowed estimation of site-specific acetylation turnover in diacetylated H3.1 (9–17) peptide, which exhibited faster apparent turnover than their monoacetylated counterparts (**Figure 3F, Supplementary Figure 4B**). Notably, we detected a trend towards higher turnover at K9 compared to K14, although the latter did not reach significance. The plateau ²H enrichment at H3K9 and H3K18 was lower than at H3K14 and H3K23, respectively (**Figure 3C, 3F**). While the mechanistic basis for these differences remains elusive, the reduced steady-state enrichment at K9 and K18 likely reflects lower stoichiometry and more rapid turnover compared to higher-occupancy sites. Alternatively, site-specific differences in acetyl-CoA sourcing may contribute to and warrant further investigation.

#### H4 (4–17) peptide

PRM analysis of diagnostic fragment ions from mono-, di-, and tri-acetylated H4 (4–17) peptides enabled site-specific quantification of acetylation turnover at K5, K8, K12, and K16 residues. As an example, **Figure 3G** shows the time-dependent increase in ²H labeling of the *y5* ion, which contains H4K16ac (**Figure 1Ciii**), allowing direct measurement of K16 acetylation turnover in mono-acetylated peptide (**Figure 3H, gray line**). K16 turnover increased progressively with higher acetylation states, consistent with enhanced acetyl-group dynamics in poly-acetylated H4 species^30^ (**Figure 3H, Supplementary Figure 4C-D**). H4K5ac turnover was directly quantified from the *b3* ion of the tri-acetylated peptide (0.32 ± 0.11 h⁻¹) (**Figure 3H, green solid line**). The same strategy was used to estimate composite turnover rates for K12/K16 and K8/K12/K16 in di- and tri-acetylated peptides using the *y7-y9* and *y12* ions, respectively (**Supplementary Figure 4E-F**). A similar fragment-ion approach was used to quantify turnover at K8 and K12 in di- and tri-acetylated species (**Figure 3I)**. However, because site-specific fragment ions were not available for these residues, turnover was inferred from differential ²H enrichment of positional fragment ions represented in the **Supplementary Figure 4E-F**. Specifically, K8 turnover was calculated from the enrichment difference between *y12* and *y7* ions of the tri-acetylated peptide, whereas K12 turnover was derived from the difference between the *y7* and *y5* ions, and between the *y9* and *y4* or *y5* ions in di-acetylated species (**Figure 3I)**. Across all sites, di- and tri-acetylated species exhibited higher turnover rates than their mono-acetylated counterparts, indicating more rapid acetyl-group exchange in poly-acetylated H4 (**Table 1**).

### Effect of Fasting on Acetylation Dynamics

To test the utility of ²H2O-based acetylation turnover approach, we compared a high-carbohydrate diet (**HCD**) with fasting, two contrasting conditions in which glucose and fatty acids are the predominating hepatic mitochondrial acetyl-CoA sources, respectively. After dietary adaptation, chow-fed mice were fasted for 48 h with free access to water. Prior to fasting, body weight did not differ between groups (**Supplementary Figure 5A**, *p* = 0.43). As expected, fasting induced a significant reduction in body weight that remained below the harmful 20% threshold (**Supplementary Figure 5A**, *p* < 0.001), attributable primarily to loss of fat mass (**Supplementary Figure 5B**, *p* = 0.001), while lean mass was significantly higher in fasted mice (**Supplementary Figure 5B**, *p* = 0.01). Liver weight was reduced by prolonged fasting (**Supplementary Figure 5C,** *p* = 0.01), but the liver-to-body weight ratio was unchanged (**Supplementary Figure 5D**), indicating that liver mass declined proportionally with overall body weight rather than reflecting a selective loss of hepatic tissue. Consistent with the absence of dietary glucose, fasting significantly reduced the respiratory exchange ratio (VCO₂/VO₂) across both light and dark cycles, reflecting a metabolic shift toward predominant fatty acid oxidation (**Supplementary Figure 5E**, *p* < 0.01) and markedly lowered blood glucose levels compared with HCD-fed mice (**Supplementary Figure 5F**, *p* < 0.01). As expected, fasting increased hepatic β-hydroxybutyrate (BHB) (**Supplementary Figure 5G**, *p* = 0.02) and 2-hydroxybutyrate (2HB) (**Supplementary Figure 5H**, p < 0.01). Elevated BHB indicates enhanced fatty acid β-oxidation and ketogenesis and a higher mitochondrial NADH/NAD⁺ ratio^75^, while increased 2-HB serves as a marker of cytosolic redox stress^76^.

To assess the effect of complete withdrawal of food for 48 h on hepatic protein acetylation, we first evaluated global lysine acetylation by lysine acetyl-specific immunoblotting. Fasting increased acetylation of hepatic non-histone proteins within the 20–250 kDa molecular weight range while modestly reducing acetylation of low-molecular-weight proteins (10–15 kDa), a region enriched in histones, compared with HCD-fed mice (**Figure 4A**). This differential acetylation pattern is consistent with fasting-induced metabolic reprogramming, whereby enhanced fatty acid oxidation and ketogenesis promote mitochondrial protein hyperacetylation, while reduced nucleocytosolic acetyl-CoA availability and altered NAD⁺-dependent deacetylase activity limit histone acetylation ^77^. To determine whether these changes extended to chromatin, acid-extracted nuclear histones were analyzed by mass spectrometry following derivatization and tryptic digestion. Across HCD and fasted livers, we identified 35 and 33 histone variants, respectively, including all core histones (H2A, H2B, H3, and H4) and linker histone H1 isoforms, with 30 variants shared between groups **(Supplementary Figure 6A**). These variants were represented by 1016 and 998 peptides in HCD and fasting conditions, respectively, of which 808 peptides were common **(Supplementary Figure 6B)**. In total, 148 acetylated histone peptides were detected in HCD-fed mice and 137 in fasted mice, with 93 acetylated peptides shared between groups (**Supplementary Figure 6C, Supplementary Table 5A&B**).

**Figure 4.**
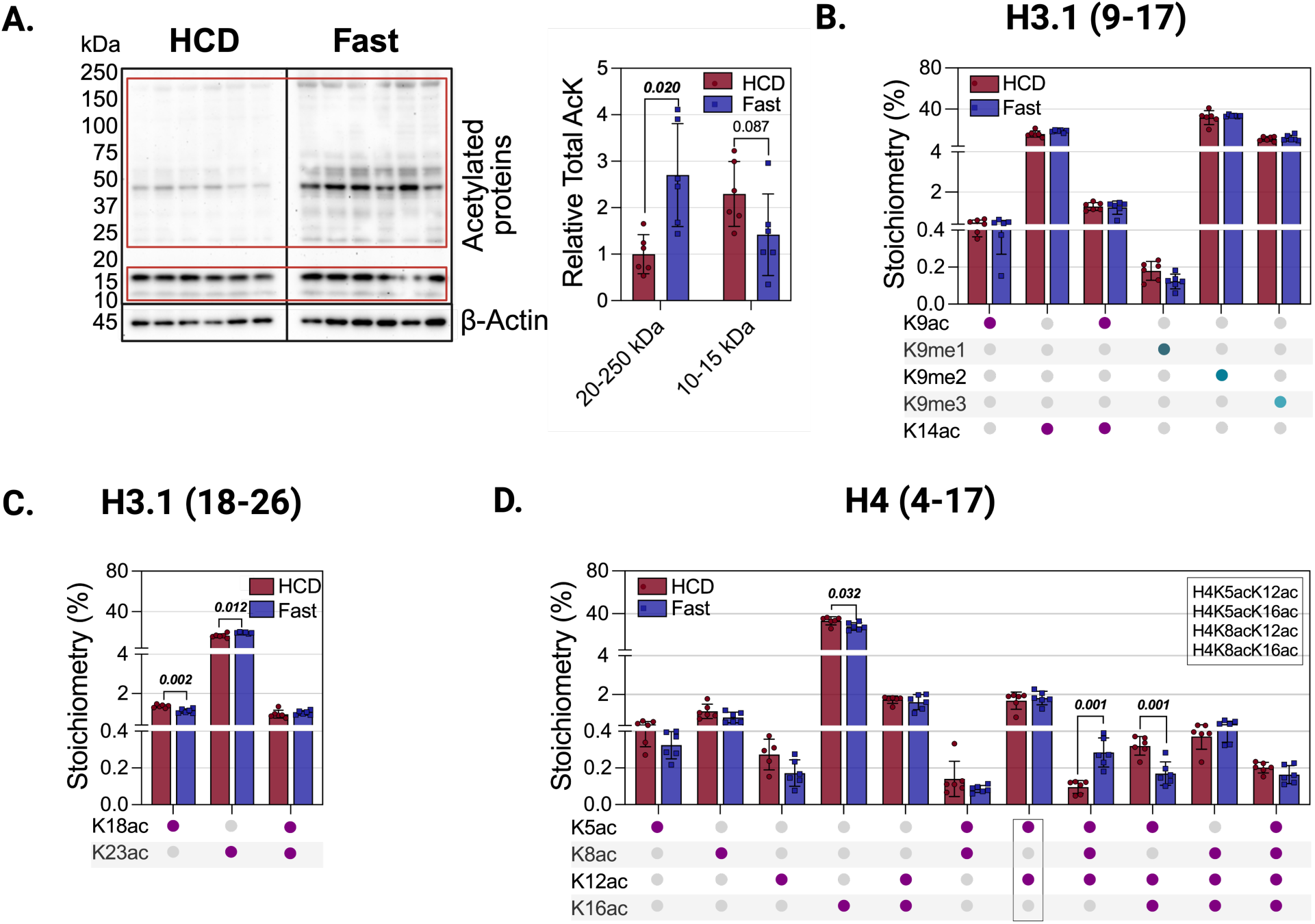
Diet-dependent changes in global protein acetylation and histone acetylation stoichiometry in mouse liver. **(A)** Global lysine acetylation of hepatic proteins in wild-type mice. Lysine-acetylated proteins from liver homogenates of high-carbohydrate diet (HCD)–fed and fasted (Fast) mice were enriched and detected by immunoblotting using a pan–acetyl-lysine antibody. Representative gels are shown, and band intensities were quantified by densitometry using AlphaView software. Normalized intensities for biological replicates (n = 6) are presented as bar graphs with error bars indicating standard deviation (SD). Due to the distinct acetylation patterns observed in low- and high-molecular-weight regions, bands within the 10–15 kDa and 20–250 kDa molecular weight ranges were quantified separately. Quantitative analysis demonstrated that fasting increased acetylation of higher-molecular-weight non-histone proteins (20–250 kDa) while modestly reduced acetylation of proteins within the 10–15 kDa range, which is enriched in core histones. (**B–C**) Site-specific acetylation stoichiometry of histone H3.1 peptides determined by LC–MS analysis. **(B)** H3.1 peptide **⁹KSTGGKAPR¹⁷** containing lysine residues K9 and K14 is mono- and di-acetylated at K9 and, mono-, di-, and tri-methylated at K9. **(C)** H3.1 peptide **¹⁸KQLATKAAR²⁶** containing lysine residues K18 and K23. Relative abundances of mono- and di-acetylated forms were used to estimate site-specific acetylation stoichiometry. **(D)** Acetylation stoichiometry of histone H4 peptide **⁴GKGGKGLGGAKR¹⁷** containing lysine residues K5, K8, K12, and K16. Acetylation states were resolved based on MS2-derived acetylation patterns. Acetylation forms that could not be distinguished among positional isomers (K5acK12ac, K5acK16ac, K8acK12ac, and K8acK16ac) are indicated in the acetylation code panel. Statistical comparisons between HCD and FAST groups were performed using two-sample *t*-tests (n = 6 per group). Relative expression levels are reported as mean ± SD, with individual data points overlaid.

Mass spectrometry–based PTM stoichiometry analysis revealed a significant increase in H3K23ac (*p* = 0.01) in fasting mice, accompanied by a reduction in H3K18ac (*p* < 0.005) (**Figure 4B–C**). In contrast, fasting significantly reduced H4K16ac (*p* < 0.05), a highly abundant site that plays a key role in regulating chromatin structure (**Figure 4D**). The selective reduction of H4K16ac during fasting is consistent with redistribution of acetyl groups toward other histones and non-histone proteins under energy-deprived conditions^78^. Collectively, these findings indicate that fasting promotes selective remodeling of histone acetylation, characterized by reduced global H4K16ac and site-specific remodeling of H3 acetylation.

Histone acetylation is regulated by acetyl-CoA availability, acetyl-CoA–dependent lysine acetyltransferases (KATs), and histone deacetylases (HDACs). Glucose-derived acetyl-CoA is generated through the sequential actions of glycolysis and the mitochondrial pyruvate dehydrogenase (PDH) complex, which converts pyruvate to acetyl-CoA. Citrate produced from this acetyl-CoA enters the cytosol and nucleus, where ATP-citrate lyase (ACLY) regenerates acetyl-CoA to support histone acetylation. In addition, acyl-CoA synthetase short-chain family member 2 (ACSS2) generates acetyl-CoA from acetate, providing an alternative substrate source for nuclear histone acetylation. Carnitine acetyltransferase (CrAT) buffers excess mitochondrial acetyl-CoA and has recently emerged as an additional regulator of acetyl-CoA availability for chromatin modification^24, 43^ (**Figure 5A**). MOF/KAT8 is the principal KAT responsible for H4K16 acetylation, and contributes to acetylation at H4K5, H4K8, H3K14, and H3K23. The NAD⁺-dependent deacetylase SIRT1 modulates H3 and H4 acetylation in a locus- and context-dependent manner during metabolic stress and aging^79^, whereas chromatin-bound SIRT6 preferentially deacetylates H3K9ac to repress glycolytic and lipogenic gene programs^80^. Under metabolic stress, cytosolic SIRT2 can transiently translocate to the nucleus and deacetylate histone substrates including H3K18ac and H4K16ac^81^.

**Figure 5.**
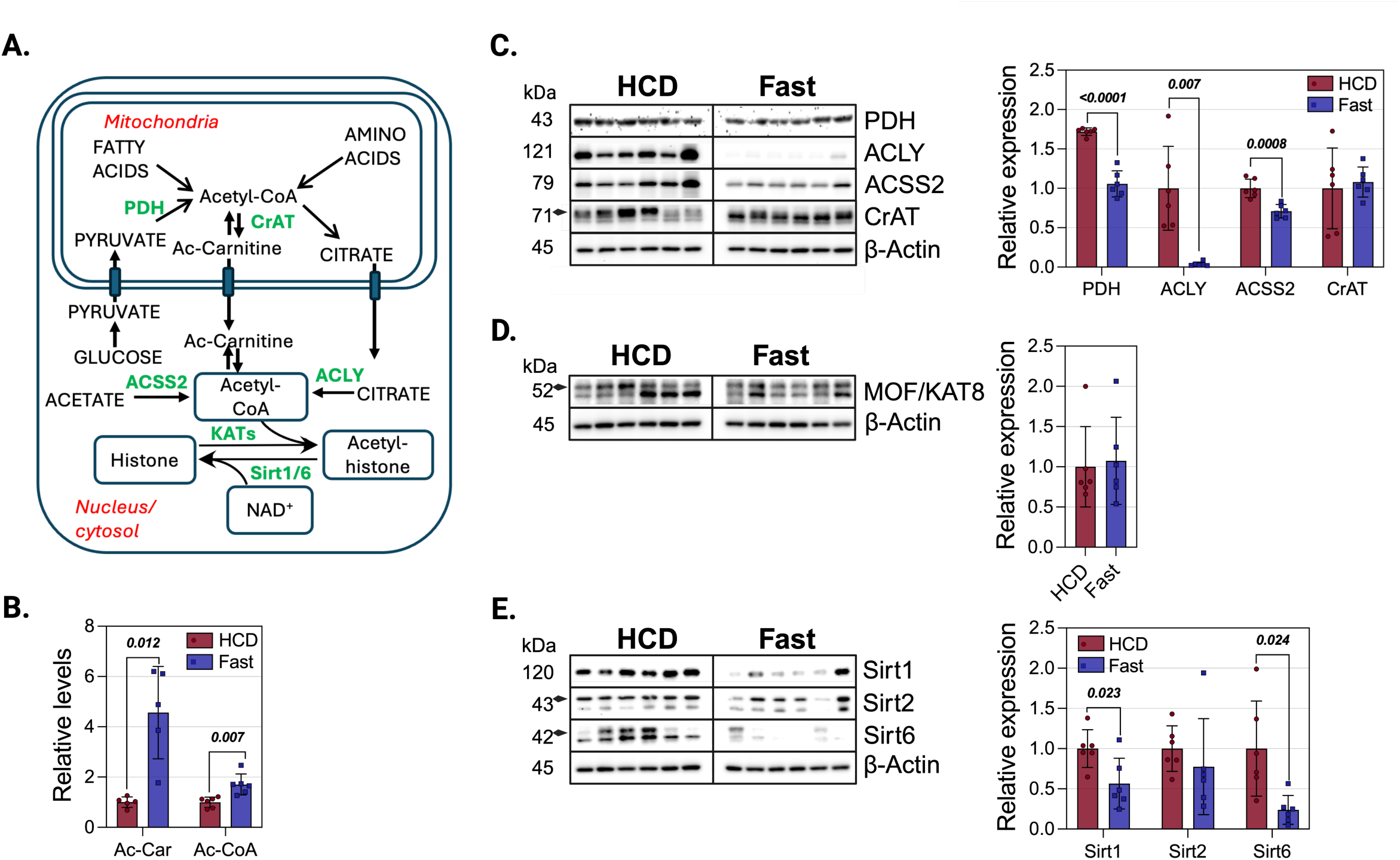
Mechanisms underlying diet-induced changes in histone acetylation in mouse liver. **(A)** Schematic overview of histone acetylation and deacetylation pathways. Histone acetylation is catalyzed by lysine acetyltransferases (KATs) using acetyl-CoA as the acetyl donor, whereas NAD⁺-dependent sirtuins catalyze histone deacetylation. The upper panel summarizes major hepatic sources of acetyl-CoA contributing to histone acetylation. **(B)** Hepatic levels of acetyl-carnitine and acetyl-CoA in high-carbohydrate diet (HCD)–fed and fasted mice. **(C)** Immunoblot analysis of enzymes regulating acetyl-CoA metabolism, including carnitine acetyltransferase (CrAT), ATP-citrate lyase (ACLY), pyruvate dehydrogenase subunit E1 (PDH E1) and acetyl-CoA synthetase 2 (ACSS2), in livers from HCD-fed and fasted mice. **(D)** Immunoblot analysis of KAT, MOF/KAT8 in livers of HCD-fed and fasted mice. **(E)** Immunoblot analysis of NAD⁺-dependent sirtuins (SIRT1, SIRT2, and SIRT6) in mouse liver under HCD and fasting conditions. **D-E**: Statistical comparisons between HCD and FAST groups were performed using two-sample *t*-tests (n = 6 per group). Relative expression levels are reported as mean ± SD, with individual data points overlaid. **C–E.** Arrowheads indicate the bands selected for quantification in the presence of non-specific signals. **Abbreviations:** ACLY, ATP-citrate lyase; ACSS2, acyl-CoA synthetase short-chain family member 2 (acetyl-CoA synthetase 2); CrAT, carnitine acetyltransferase; KAT, lysine acetyltransferase; PDH, pyruvate dehydrogenase; SIRT1/6, sirtuin 1 and sirtuin 6.

Fasting increased hepatic acetyl-carnitine approximately fourfold compared with the HCD group (**Figure 5B**, *p* < 0.05), consistent with enhanced mitochondrial fatty-acid β-oxidation and elevated acetyl-CoA production. Excess mitochondrial acetyl-CoA is buffered by CrAT, which converts acetyl-CoA to acetyl-carnitine to maintain the free CoA pool. Although fasting also increased total hepatic acetyl-CoA, the change was approximately twofold smaller than that observed for acetyl-carnitine (**Figure 5B**, *p* = 0.007), reflecting rapid utilization of acetyl-CoA through ketogenesis and tricarboxylic acid (TCA) cycle oxidation. These results indicate that acetyl-carnitine primarily reflects mitochondrial acetyl-CoA overflow rather than total cellular acetyl-CoA abundance. Accordingly, carbohydrate feeding favors nucleo-cytosolic PDH-dependent acetyl-CoA production that supports ACLY–regulated lipogenesis and histone acetylation, whereas fasting shifts metabolism toward mitochondrial oxidation and ketone production, thereby limiting substrate availability for anabolic and epigenetic processes.

Consistent with reduced glucose-derived acetyl-CoA production, 48-h fasting significantly decreased hepatic PDH E1 subunit, nearly abolished ACLY and significantly decreased ACSS2 expression, while CrAT levels remained unchanged (**Figure 5C).** These findings indicate a marked suppression of glycolytic and acetate-dependent nucleo-cytosolic acetyl-CoA generation pathways while preserving mitochondrial acetyl-CoA buffering capacity. Notably, MOF/KAT8 protein abundance remained unchanged despite a marked reduction in H4K16ac (**Figure 5D**), suggesting that fasting-induced depletion of the nucleo-cytosolic acetyl-CoA pool, rather than reduced MOF expression, contributes to decreased H4K16 acetylation.

Fasting also significantly reduced hepatic SIRT1 and nearly abolished SIRT6, while SIRT2 remained stable (**Figure 5E**). Despite loss of SIRT6, bulk nuclear H3K9ac was unchanged (**Figure 4B**), indicating that steady-state H3K9 acetylation is buffered by reduced acetylation input and/or compensatory class I/II HDAC activity rather than being determined solely by SIRT6 expression. In contrast, decreased H3K18ac despite reduced SIRT1 suggests that H3K18 acetylation is primarily limited by diminished acetyl-CoA availability rather than enhanced sirtuin-mediated deacetylation. Together, these findings suggest that prolonged fasting shifts the hepatic chromatin landscape predominantly through changes in acetylation flux and enzymatic activity at specific loci, rather than uniform regulation of sirtuin protein abundance.

Next, we tested the sensitivity of the ²H₂O labeling approach described above to detect diet-induced changes in acetylation turnover by comparing fasting and HCD-fed mice. Following a bolus injection of ²H₂O-saline, mice from both HCD-fed and fasted groups were maintained on 8% ²H₂O in drinking water for up to 24 h. Body-water enrichment stabilized within 1 h at 3.8 ± 0.4% in HCD-fed mice and 3.5 ± 0.3% in fasted mice (**Supplementary Figure 7A**). Glucose ²H enrichment in fasted mice rapidly plateaued at ∼12% within 2 h, whereas in HCD-fed mice enrichment was slightly lower (∼7–8%) (**Supplementary Figure 7B**). In contrast, palmitate showed minimal (∼1%) ²H enrichment in fasted mice, indicating negligible de novo lipogenesis, but increased linearly to ∼10% in HCD-fed mice (**Supplementary Figure 7C)**. The minmal ²H enrichment of palmitate in fasted mice indicates that fatty-acid–derived acetyl-CoA remained largely unlabeled under these conditions.

Consistent with rapid acetyl-group exchange, ²H₂O labeling revealed that acetyl-carnitine turned over extremely rapidly (*t½* ≈ 0.4 h) in both dietary states, with no significant difference in plateau ²H enrichment (**Supplementary Figure 7D**), reflecting dominant CrAT-mediated reversible buffering of mitochondrial acetyl groups. In contrast, total tissue acetyl-CoA exhibited marked diet sensitivity: fasting increased acetyl-CoA turnover by >3-fold while suppressing plateau ²H enrichment by >30% (**Supplementary Figure 7E**), indicating accelerated acetyl-CoA flux coupled to dilution by unlabeled acetyl units derived predominantly from fatty-acid β-oxidation. Thus, these findings indicate that acetyl-CoA labeling captures diet-dependent shifts in carbon sources and metabolic flux. Accordingly, carbohydrate feeding favors ACLY-supported nucleo-cytosolic acetyl-CoA availability for lipogenesis and histone acetylation, whereas fasting reallocates acetyl-CoA toward mitochondrial oxidation and ketone production.

Full-scan MS1 spectra demonstrated progressive ²H incorporation into acetylated H3.1 9–17, H3.1 18–26 (**Figure 6A–D**), and H4 4–17 (**Figure 6E–H**) peptides in both HCD and fasted mouse livers (**Table 2**). No significant differences in acetylation turnover were observed for mixed mono-acetylated H3.1 (18–26) (**Figure 6A**) or H3.1 (9–17) peptides (**Figure 6C**). In contrast, turnover of mixed mono-acetylated H4 (4–17) peptides was significantly increased in fasted mice (*p* < 0.05; **Figure 6E**). Consistently, multi-acetylated H3.1 (9–17) (**Figure 6D**) and H4 (4–17) (**Figure 6E–F**) peptides also exhibited faster turnover under fasting conditions. The di-acetylated H3.1 (18-26) (**Figure 6D**) and tetra-acetylated H4 (4-17) peptides (**Figure 6H**) showed a similar upward trend, although this did not reach statistical significance, likely due to greater variability associated with low stoichiometry and limited abundance. As anticipated, plateau ²H enrichment of acetylated peptides was lower during fasting, likely reflecting dilution of labeled acetyl-CoA by unlabeled acetyl-CoA generated from fatty acid β-oxidation. These results are consistent with prior studies demonstrating that, under glucose-limited conditions, lipids become a predominant carbon source for histone acetylation, thereby contributing to metabolic reprogramming^82^.

**Figure 6.**
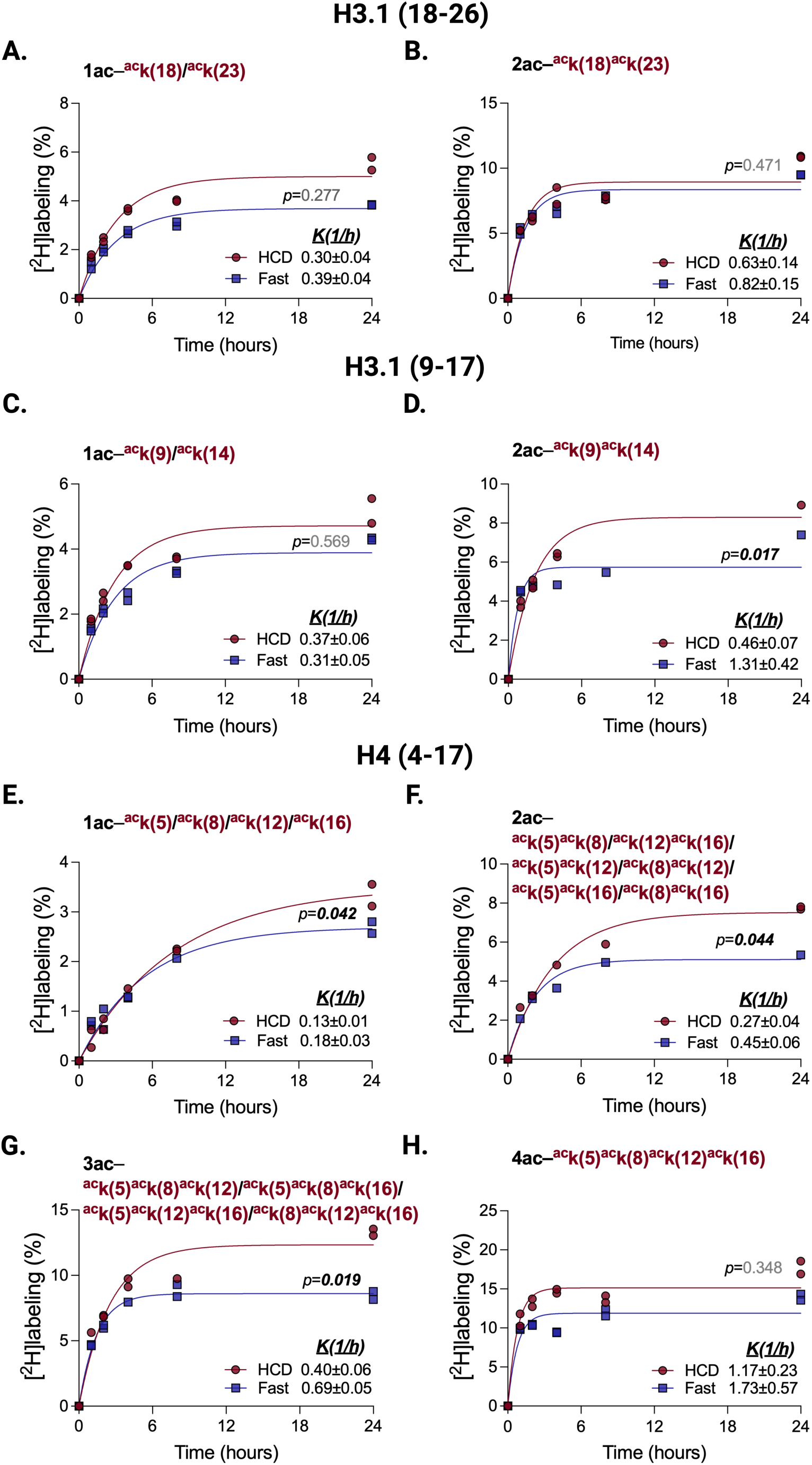
MS1-based quantification of diet-dependent changes in histone acetylation turnover and acetyl-CoA sources in vivo. Time-dependent incorporation of ²H into acetylated histone peptides was determined from MS1 isotopomer distributions of mono- and di-acetylated H3.1 (18–26) peptides (**A–B**), mono- and di-acetylated H3.1 (9–17) peptides (**C–D**), and mono- (**E**), di- (**F**), tri- (**G**), and tetra-acetylated (**H**) H4 (4–17) peptides. Increasing numbers of neighboring acetylation sites were associated with proportionally higher acetylation turnover under both dietary conditions. Fasting increased acetylation turnover while reducing the plateau ²H enrichment of the acetyl group, indicating a greater contribution of unlabeled acetyl-CoA derived from fatty-acid β-oxidation to histone acetylation. Statistical comparisons between HCD and FAST groups were performed using *F*-tests (n = 6 per group). Turnover rate estimates are reported as mean ± SE.

**Table 2.**
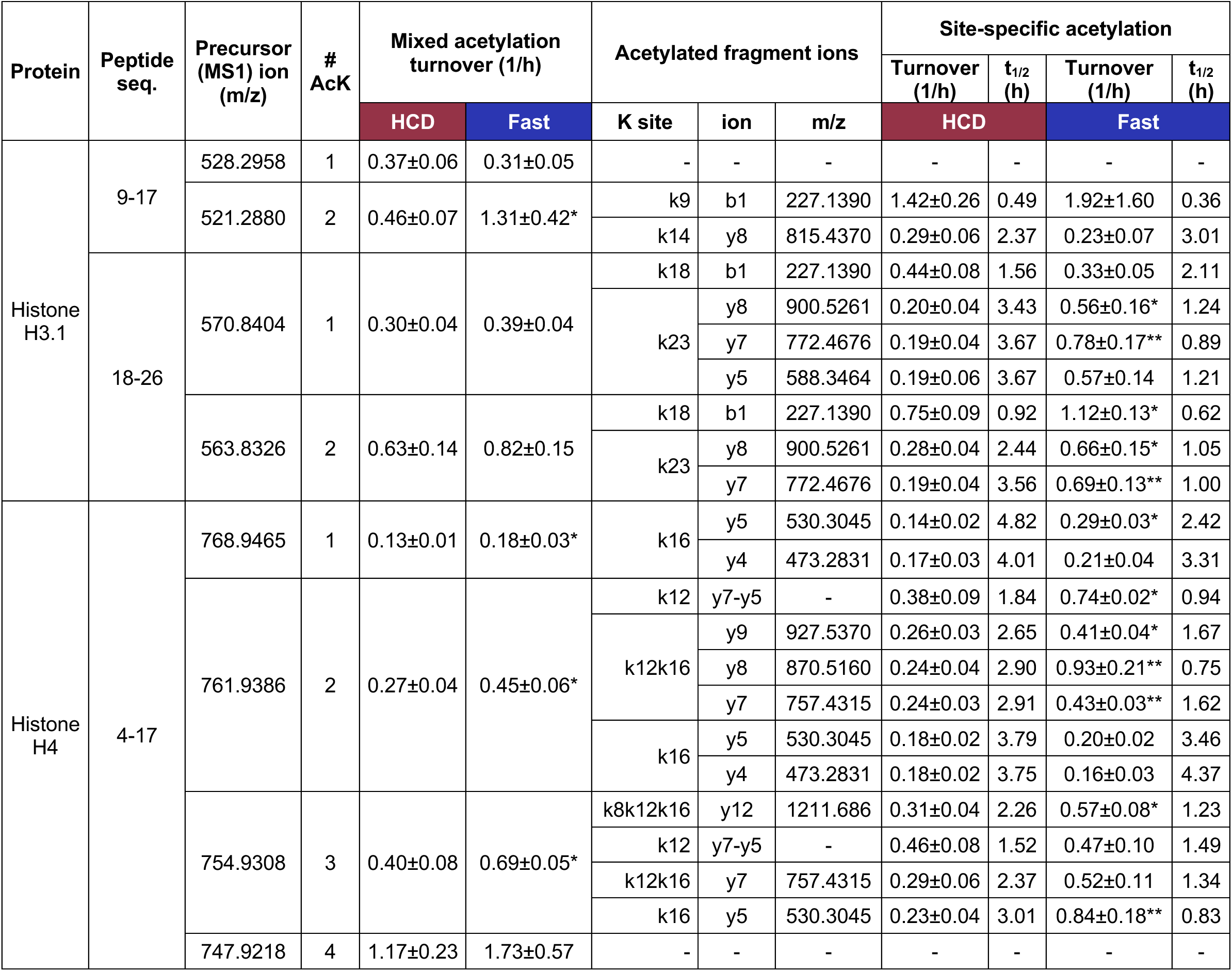
Site-specific histone acetylation turnover in HCD-fed and fasted mouse liver quantified by fragment-ion–resolved ²H. ₂**O metabolic labeling.** The table summarizes acetylated histone peptides, the precursor and diagnostic fragment ions used for composite and site-specific analyses, and the corresponding acetylation turnover rates determined from time-dependent ²H incorporation into precursor and fragment-ion isotopomers. Turnover parameters are reported for mice maintained on a high-carbohydrate diet (HCD) or subjected to 48-h fasting. *p < 0.05 and **p < 0.005.

Although MS1-based quantification of total ^2^H labeling in acetylated peptides provides a robust measure of global acetylation dynamics and demonstrates that short-term (24 h) ^2^H2O metabolic labeling can reveal the relative contribution of non-glycolytic substrates (e.g., fatty acids) to histone acetylation, resolution of turnover kinetics at individual lysine residues requires analysis of site-isolated acetylated species. To resolve site-specific acetylation dynamics within these peptides, we next applied targeted PRM fragment-ion analysis. High-resolution MS/MS analysis of diagnostic *b* and *y* fragment ions enabled site-specific quantification of acetylation turnover at H3.1 K9 and K14 within the H3.1 (9–17) peptide and at K18 and K23 within the H3.1 (18–26) peptide. Consistent with MS1-based measurements, isotope distribution of high-resolution b1 and y7 fragment ions showed that fasting significantly increased acetylation turnover at K18 and K23 in the di-acetylated H3.1 (18–26) peptide (**Figure 7Aiii**), as well as at K23 in the mono-acetylated (18–26) peptide (**Figure 7Aii),** without any significant change in K18 acetylation turnover (**Figure 7Ai**). Whereas acetylation turnover at K9 and K14 in the low-abundance di-acetylated H3.1 (9–17) peptide was unaffected by fasting (**Figure 7Bi–ii; Supplementary Figure 8C; Table 2**). These findings were confirmed by analysis of the *y5* and *y8* fragment ions (**Supplementary Figure 8A-C**).

**Figure 7.**
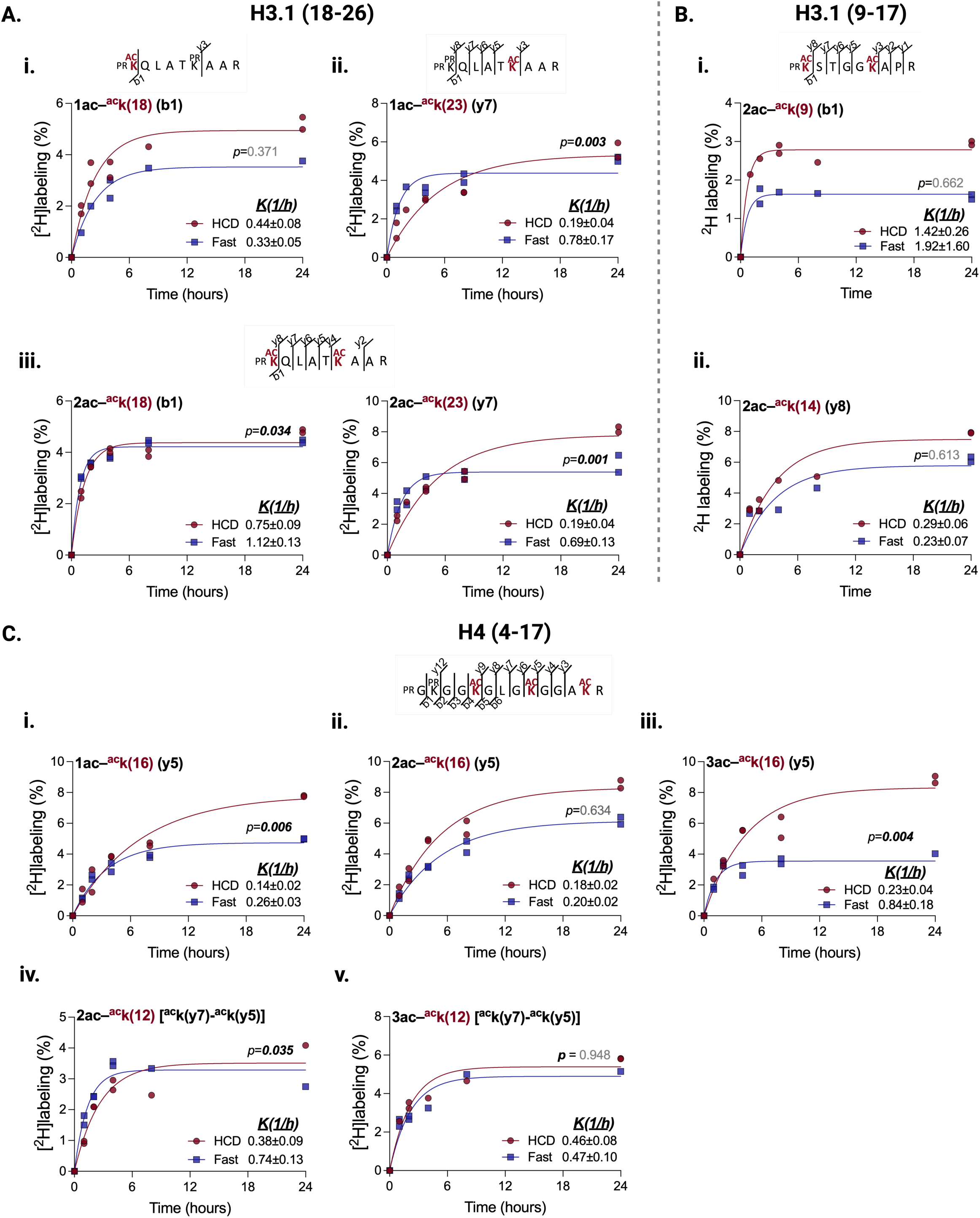
MS2 (fragment-ion)–based quantification of diet-dependent, site-specific histone acetylation turnover in vivo. ²H incorporation into fragment ions containing the acetyl group was quantified for mono- and di-acetylated H3.1 (18–26) peptides (**A**), di-acetylated H3.1 (9–17) peptides (**B**), and mono-, di-, and tri-acetylated H4 (4–17) peptides (**C**). **(A)** Time-dependent ²H incorporation into diagnostic ***b1*** and ***y7*** fragment ions enables site-specific acetylation turnover measurements in mono- (**i–ii**) and di-acetylated (**iii**) H3.1 (18–26) peptides. **(B)** Time-dependent ²H incorporation into diagnostic ***b1*** and ***y8*** fragment ions enables site-specific turnover measurements in di-acetylated (**i–ii**) H3.1 (9–17) peptides. **(C)** Time-dependent ²H incorporation into diagnostic ***y5*** fragment ions from mono- (**i**), di- (**ii**), and tri-acetylated (**iii**) H4 (4–17) peptides enables quantification of acetylation turnover at H4K16. Site-specific acetylation turnover at H4K12 was estimated from calculated ²H enrichment using ***y5*** and ***y7*** fragment ions in di- (**iv**) and tri-acetylated (**v**) H4 (4–17) peptides. Increased labeling with higher acetylation states reflects enhanced acetylation turnover in polyacetylated H4 peptides.

Similarly, the fragment ion analysis demonstrated that fasting increased acetylation turnover at lysine 16, in mono- and tri-acetylated H4 4-17 peptides (**Figure 7Ci, 7Ciii, Supplementary Figure 9C)** without any differences in diacetylated peptide (**Figure 7Cii, Supplementary Figure 9B**). In the mono-acetylated peptide, significance was observed for the *y5* ion, whereas the *y4* ion exhibited a similar but nonsignificant increase (**Supplementary Figure 9A)**. We also estimated acetylation turnover at lysine 12 based on its calculated ^2^H-enrichment using *y7* and *y5* fragment ions of di- and tri-acetylated species. While fasting was associated with doubled turnover rate at lysine 12 of diacetylated peptide (**Figure 7Civ)**, no significant change was observed in tri-acetylated peptide (**Figure 7Cv)**. Notably, analysis of the *y7, y8, and y9* fragment ions in the di- and tri-acetylated peptides also indicated increased mixed K12/K16 acetylation turnover under fasting conditions (**Supplementary Figure 9C–E**). In addition, y7 and *y12* fragment-ion analysis of the tri-acetylated H4 (4–17) peptide revealed an approximately twofold increase in combined acetylation turnover at K12, and K16, and, K8, K12, and K16 residues, respectively (**Supplementary Figure 9F-G, Table 2**).

To assess whether site-specific acetylation turnover depends on acetylation abundance, we performed Pearson correlation analysis between pooled site-specific turnover rates and stoichiometry. A negative trend was observed (r = −0.299), indicating that more highly occupied sites tended to exhibit slower turnover, although the correlation did not reach statistical significance (*p* = 0.096) (**Supplementary Figure 10**).

Fragment-ion–resolved analysis also confirmed that the ²H₂O labeling approach can detect shifts in the metabolic origin of acetyl-CoA used for histone acetylation. Using plateau ²H enrichment of acetylated peptides and kinetic modeling, we previously estimated the maximum number of exchangeable hydrogens within the acetyl group in mouse liver^83^, demonstrating incorporation of ∼2.75 deuterium atoms under fed conditions where glucose predominates as an acetyl-CoA source^77^. Fasting reduced plateau ²H enrichment of acetyl groups at K18 and K23 in mono-acetylated H3.1 (18–26) peptides (**Figure 8A**) and at K9 in the di-acetylated H3.1 (9–17) peptide (**Figure 8B**), consistent with increased incorporation of fatty acid–derived (“cold”) acetyl-CoA during fasting. A similar decrease in plateau enrichment was observed for the high-occupancy K16-containing mono-, di-, and tri-acetylated H4 (4–17) peptides (**Figure 8C)**. However, accurate determination of plateau enrichment in the di-acetylated H3.1 (9–17) and (18–26) peptides, as well as in the di- and tri-acetylated H4 (4–17) species, was limited by their low stoichiometry (∼1%) (**Figure 8A-C)**. This low abundance reduced quantitative precision and limited the ability to robustly resolve fasting-induced changes in the acetyl-CoA sources contributing to these acetylation events.

**Figure 8.**
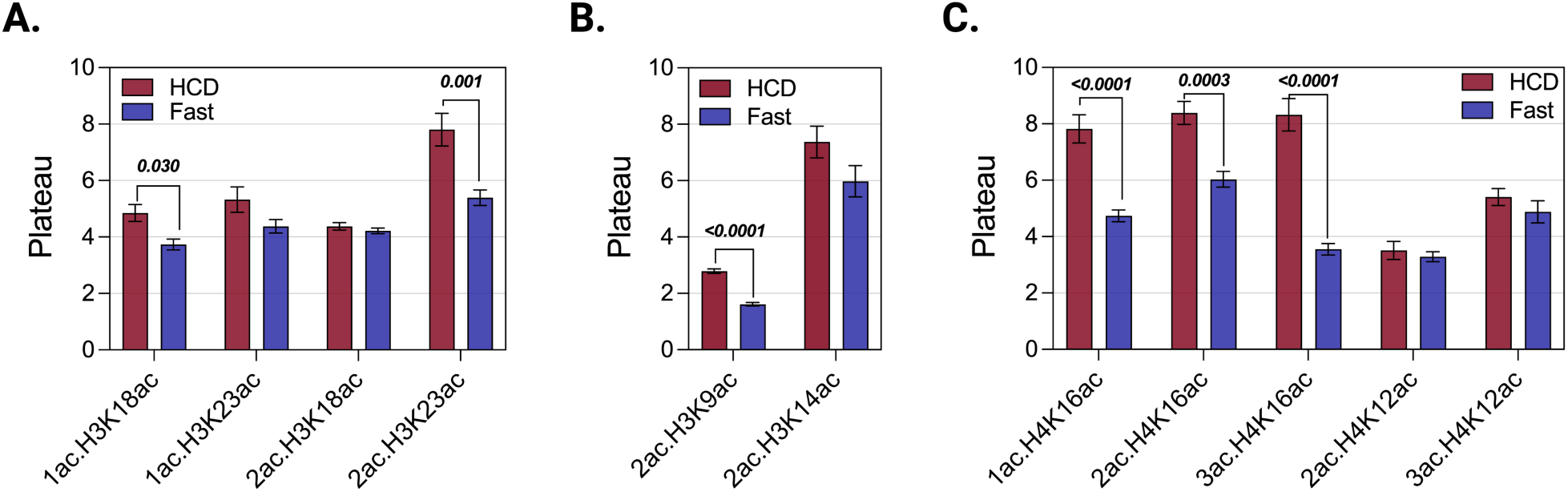
Estimated contribution of non-glycolytic acetyl-CoA to site-specific hepatic histone acetylation during fasting. Fasting-induced changes in plateau ²H enrichment of diagnostic fragment ions from acetylated histone peptides were compared with those in mice fed a high-carbohydrate diet (HCD). Reduced plateau enrichment during fasting reflects dilution of glucose-derived ²H-labeled acetyl-CoA with unlabeled (“cold”) acetyl-CoA generated from fatty-acid β-oxidation. Because fatty acids incorporate negligible ²H during short-term (<24 h) ²H₂O metabolic labeling, increased fatty-acid oxidation lowers the isotopic enrichment of the acetyl group used for histone acetylation. Statistical comparisons of plateau ^2^H-enrchment between HCD and Fast groups were performed using extra sum-of-squares *F*-test. A global model sharing a common plateau parameter was compared against an unconstrained model allowing for independent fits (n = 6 per group). Plateau estimates are reported as mean ± SE.

## CONCLUSION

We developed a stable-isotope–enabled mass spectrometry workflow that enables simultaneous quantification of histone acetylation stoichiometry, site-specific acetylation turnover, and acetyl-CoA substrate contributions *in vivo*. By combining short-term ^2^H2O metabolic labeling with high-resolution Orbitrap mass spectrometry, propionylation-based histone derivatization, and fragment-ion–resolved PRM analysis, we overcame major analytical challenges associated with chromatographically unresolved positional acetylation isomers and enabled direct measurement of acetyl-group turnover independent of histone protein turnover *in vivo*.

Application of this approach to mouse liver demonstrated that for major H3.1 and H4 acetylation sites histone acetylation turnover occurs on the scale of hours, several orders of magnitude faster than histone protein turnover. MS1-based measurements provided robust quantification of global acetylation dynamics, whereas fragment-ion analysis enabled site-specific turnover measurements for individual lysine residues within combinatorially modified peptides. Furthermore, acetyl-group plateau ²H enrichment served as a quantitative readout of the relative contributions of glycolytic and alternative substrates to the acetyl-CoA pool. Using fasting as a physiologically relevant metabolic perturbation, we demonstrate that alterations in nutrient availability remodel histone acetylation through coordinated changes in acetylation stoichiometry, acetyl-CoA metabolism, and acetylation turnover. Fasting increased turnover at selected H3.1 and H4 acetylation sites while decreasing acetyl-group labeling, indicating a shift from glucose-derived acetyl-CoA in the fed state to greater utilization of fatty acid–derived acetyl-CoA during fasting. These findings highlight the dynamic coupling between cellular metabolism and chromatin regulation and illustrate the utility of ^2^H2O labeling for interrogating epigenetic metabolic fluxes in vivo.

Several limitations should be considered. Accurate estimation of acetylation turnover rates and plateau enrichments becomes increasingly challenging for low-stoichiometry acetylation sites and multiply acetylated peptides because limited ion abundance reduces quantitative precision. In addition, the chromatographic method did not resolve positional acetylation isomers, precluding reliable MS1-based quantification and necessitating site-specific fragment ion (MS2)-based analysis. Although higher-resolution chromatographic separations or ion mobility mass spectrometry could improve isomer resolution, consistent measurements across multiple site-diagnostic fragment ions substantially mitigated this limitation. Interpretation of acetyl-group labeling also assumes isotopic and metabolic steady state throughout the labeling period; therefore, rapidly changing physiological conditions or incomplete precursor equilibration may introduce uncertainty into estimates of acetylation turnover and acetyl-CoA substrate contributions. For certain H4 acetylation sites, turnover rates were derived from differences in isotopic enrichment between complementary fragment ions rather than measured directly, potentially increasing analytical uncertainty. Furthermore, the current approach is optimized for rapidly turning over, enzymatically regulated histone acetylation, where short-term ²H₂O labeling selectively labels newly synthesized acetyl groups while peptide backbones remain largely unlabeled. Extension to slowly turning over acetylation sites, particularly non-enzymatic acetylation of long-lived non-histone proteins, will require correction for concurrent peptide backbone labeling and more comprehensive kinetic modeling. Finally, advances in instrument sensitivity, ion mobility separation, data-independent acquisition, and complementary stable-isotope tracing strategies should further improve coverage of low-abundance acetylation sites and complex histone proteoforms.

Despite these limitations, this work establishes the first integrated approach for simultaneous measurement of histone acetylation occupancy, turnover, and acetyl-CoA source utilization *in vivo*. The ability to connect chromatin modifications directly to underlying metabolic fluxes provides a powerful tool for investigating epigenetic regulation in physiology and disease. Importantly, whereas previous stable-isotope studies of histone acetylation turnover have been largely restricted to cultured cells using ¹³C-labeled substrates, and no method has been available for direct measurement of histone acetylation turnover *in vivo*, the present approach enables such measurements using inexpensive ²H₂O administered through a simple intraperitoneal priming dose followed by enriched drinking water. This minimally invasive labeling strategy simultaneously quantifies histone acetylation stoichiometry, turnover, and acetyl-CoA source utilization in freely moving animals without the need for costly tracers or complex infusion protocols. More broadly, ²H₂O labeling offers a simple, scalable, and cost-effective approach for studying metabolic control of histone acetylation *in vivo*, with applications to aging, cancer, neurodegeneration, and other disorders characterized by altered acetyl-CoA metabolism.

## Supplementary material

Supplementary methods and data analysis

Supplementary Tables S1-S5

Supplementary Figures S1-S10.

## Author contributions

Conceptualization: T.K., T.-H.T., and G.-F.Z. Methodology: A.A.-A., S.I., M.A., W.H., Formal Analysis: A.A.-A., S.I., T.-H.T., Visualization: T.-H.T., T.K., A.A.-A., Supervision: T.K. and G.-F.Z., Writing: T.K., Review and editing: A.A.-A., S.I., M.A., W.H., T.-H.T., G.-F.Z., and T.K.

## Funding

This work was supported in parts by the National Institute of Health grants R21 AA029784, R01AA030026 and 1R21AG085590-01 (T.K.), R21AG091377 (T.K.) and R21 TR005163 (G.-F.Z.).

## Acknowledgements

We thank Dr. Usman Sabir and Henyah Dardir for their technical assistance. Figures are created in BioRender. Alvarado, A. (2026) https://BioRender.com/lpdz3qz.

## Data availability

All data needed to evaluate the conclusions in the paper are present in the paper and the supplementary materials. The mass spectrometry proteomics data have been deposited to the ProteomeXchange Consortium via the PRIDE partner repository with the ProteomeXchange (accession number: PXD076407 and 10.6019/PXD076407, <u>reviewer account details for acessing the dataset: username:</u>, password: 6h5qBVP9Itep).

Annotated spectra that support the identification of acetylated peptides are available in the supplementary materials section of this submission.

## Conflict of interest

All authors declare no conflicts of interest.

## Supplementary Figures

**Supplementary Figure 1.**
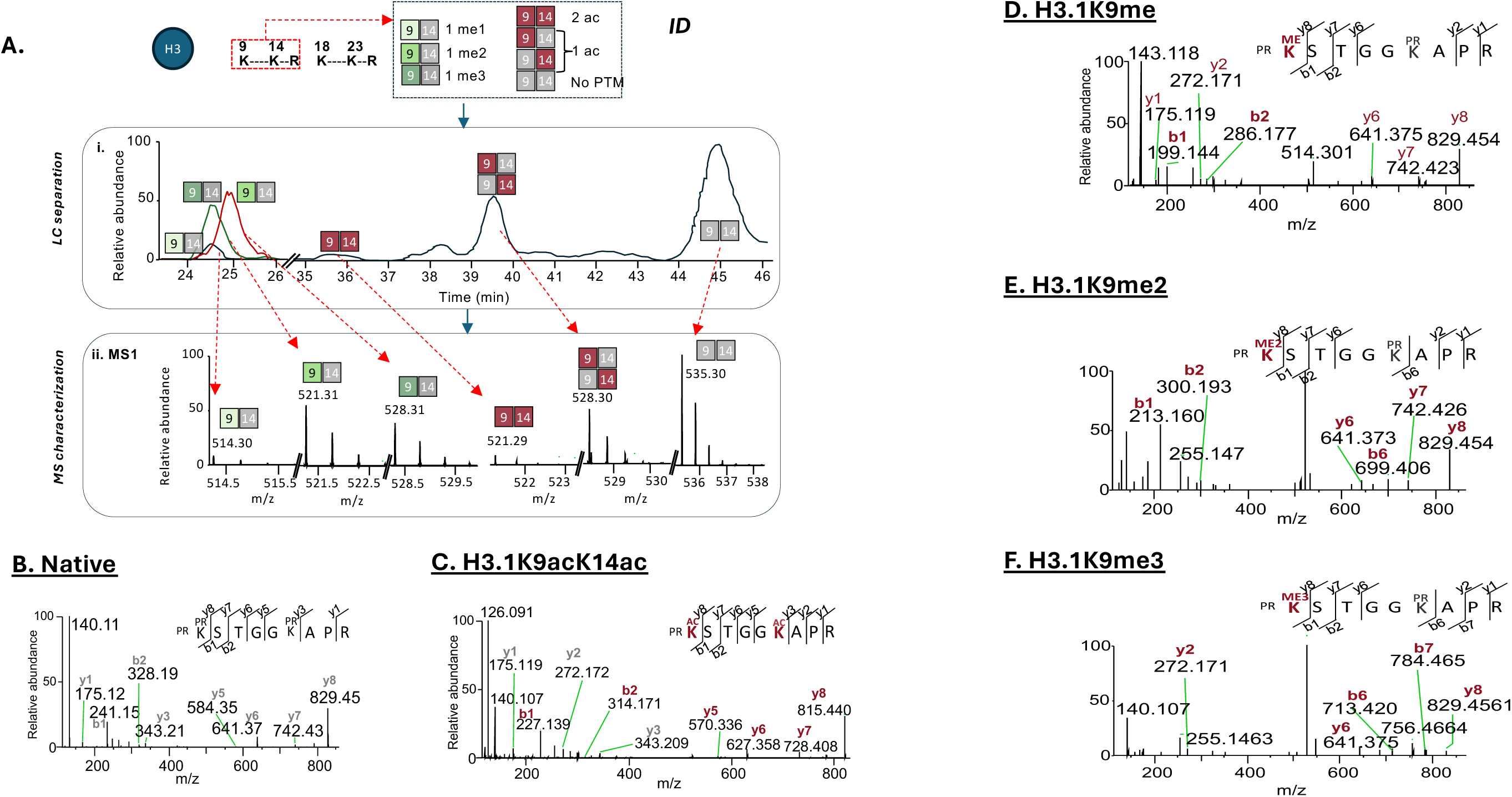
Chromatographic separation and MS characterization of site-specific methylation and acetylation of histone H3.1 peptides (residues 9–17). (A) (i) Representative chromatographic separation of H3.1 (9–17) peptides containing lysine 9 mono-, di-, and trimethylation (K9Me1, K9Me2, K9Me3), mono-acetylation at K9 or K14 (K9Ac/K14Ac), diacetylation (K9AcK14Ac), and the native chemically dipropionylated form (K9PrK14Pr). (ii) MS1-based characterization of the corresponding methylated and acetylated peptide species. (B–F) MS/MS characterization using targeted parallel reaction monitoring (PRM) of the native H3.1 (9–17) peptide (B), diacetylated K9AcK14Ac (C), K9Me1 (D), K9Me2 (E), and K9Me3 (F). Diagnostic *b* and *y* fragment ions enable discrimination of distinct methylated proteoforms and positional acetylation isomers at individual lysine residues..

**Supplementary Figure 2.**
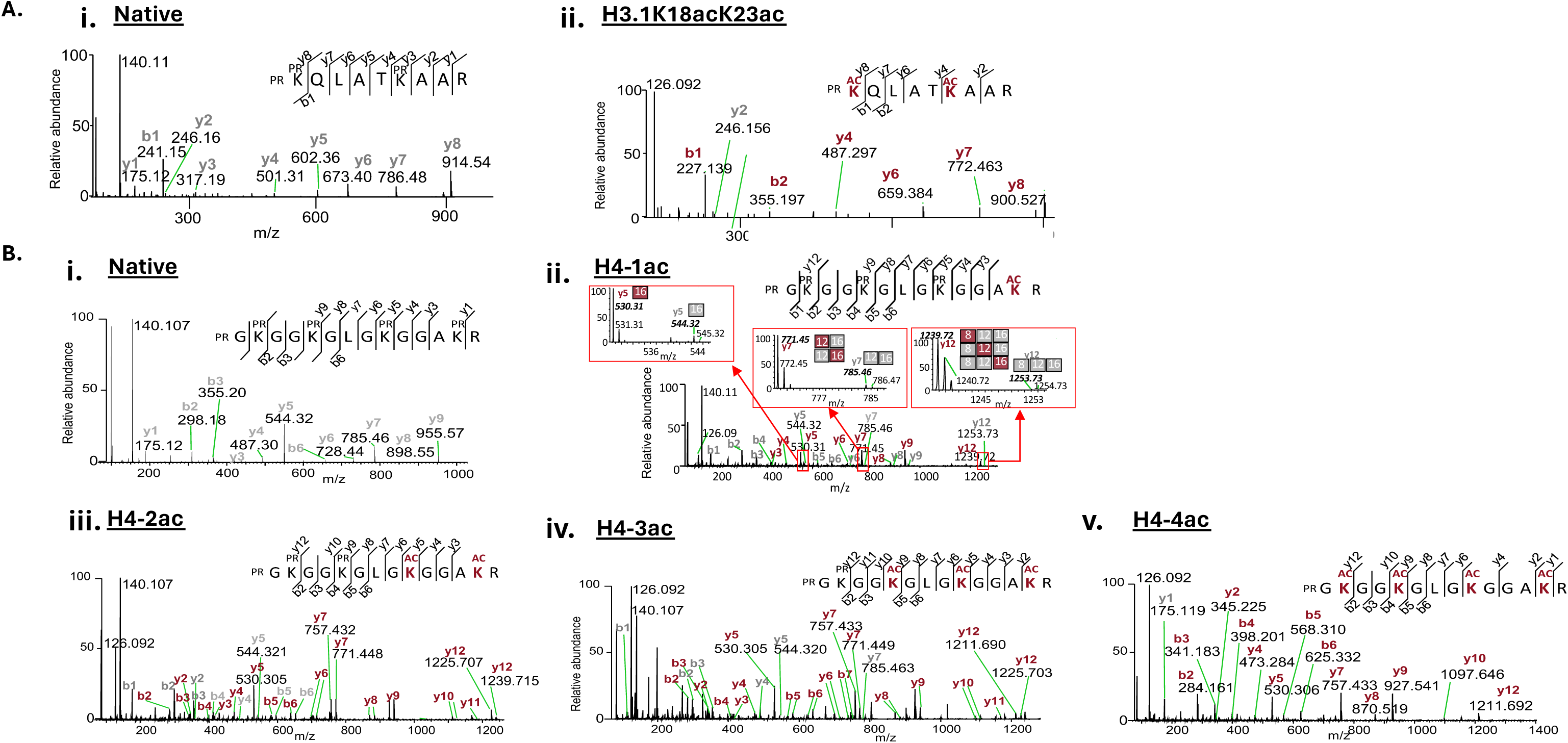
MS/MS characterization of acetylated histone H3.1 and H4 peptides by targeted PRM. Targeted parallel reaction monitoring (PRM) was used to characterize acetylated histone peptides and confirm site-specific acetylation using diagnostic fragment ions. (A) H3.1 (18–26) peptide: native propionylated form (Ai) and diacetylated K18AcK23Ac species (Aii). (b) H4 (4–17) peptide: native all propionylated form (bi), mono-acetylated K16Ac (bii), di-acetylated K12AcK16Ac (H4K12acK16ac), tri-acetylated K8AcK12AcK16Ac (biv), and tetra-acetylated K5AcK8AcK12AcK16Ac species (bv). Diagnostic *b* and *y* fragment ions enable confirmation of acetylation states and site localization at individual lysine residues.

**Supplementary Figure 3.**
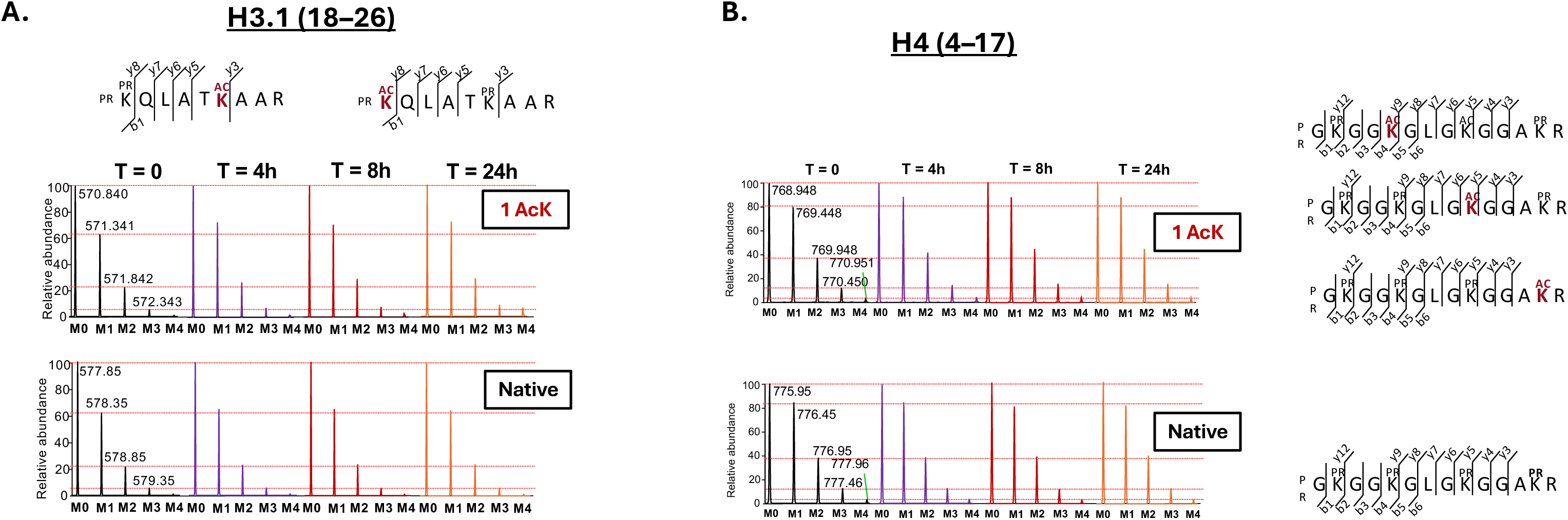
MS1 isotopomer distributions of acetylated histone peptides during ²H₂O labeling. **(A)**Representative high-resolution MS1 spectra of isobaric mono-acetylated H3.1 (18–26) peptides (K18Ac or K23Ac) and the corresponding native peptide. (**B**) Representative high-resolution MS1 spectra of mono-acetylated H4 (4–17) peptides (K5Ac, K8Ac, K12Ac, or K16Ac) and the corresponding native peptide. Progressive enrichment of M+1–M+3 isotopomers in acetylated peptides reflects incorporation of ²H from body water into the acetyl moiety, whereas the native peptides show no detectable isotopic shift.

**Supplementary Figure 4.**
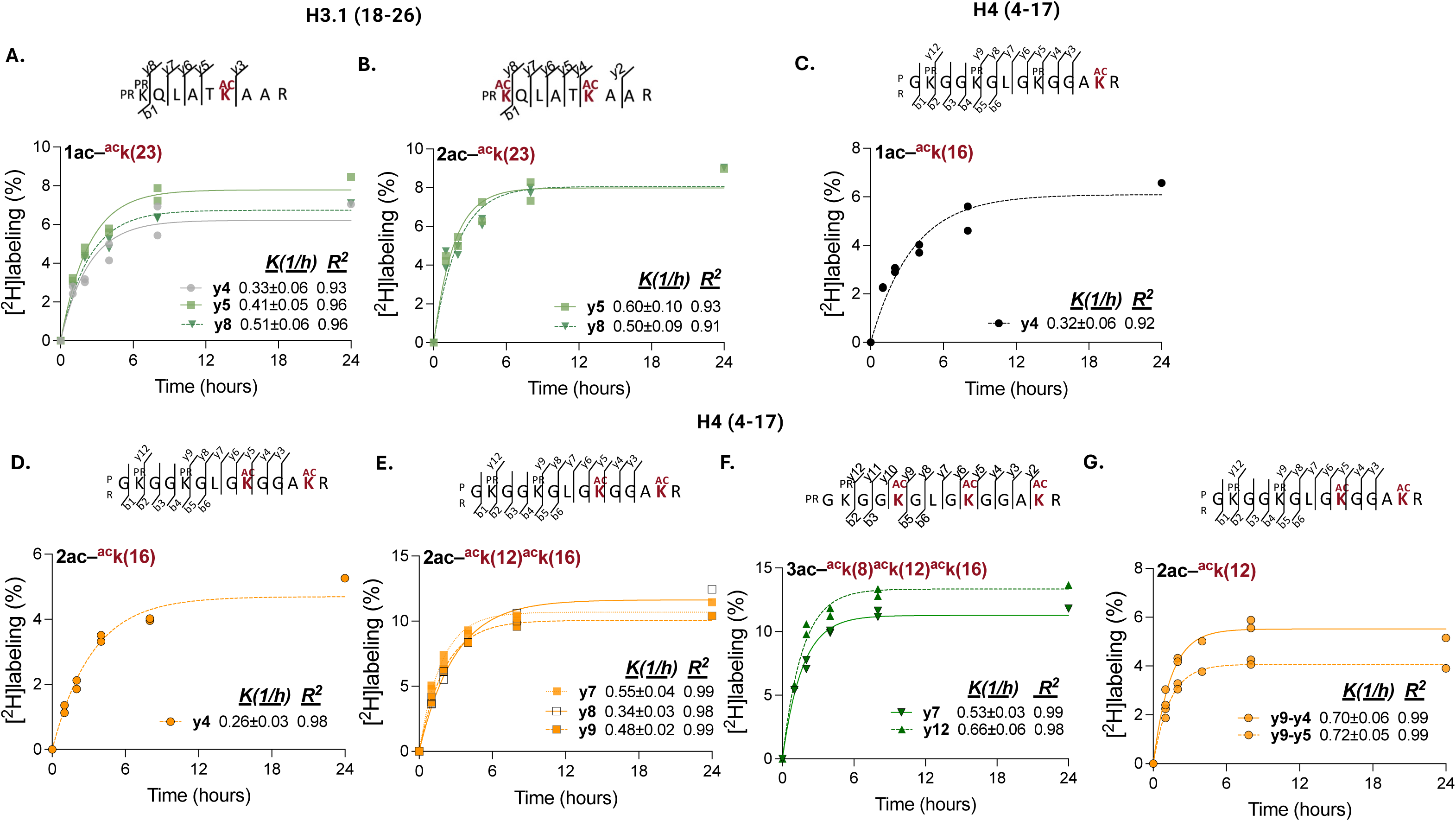
MS2 (fragment ion)–based ²H₂O metabolic labeling for quantifying site-specific histone acetylation turnover in vivo. **(A)** Time-dependent ²H incorporation into *y4, y5,* and *y8* fragment ions from mono-acetylated H3.1 (18–26) peptide enables quantification of acetylation turnover at K23. **(B)** Time-dependent ²H incorporation into *y5* and *y8* fragment ions from di-acetylated H3.1 (18–26) peptide enables quantification of acetylation turnover at K23. **(C)** Time-dependent ²H incorporation into *y4* fragment ions from mono-acetylated H4 (4–17) peptide enables quantification of acetylation turnover at K16. **(D)** Time-dependent ²H incorporation into *y4* fragment ions from di-acetylated H4 (4–17) peptide enables quantification of acetylation turnover at K16. **(E)** Time-dependent ²H incorporation into *y7-y9* fragment ions containing acetylated K12 and K16 from di-acetylated H4 (4–17) peptides enables quantification of composite acetylation turnover at H4K12 and H4K16. **(F)** Time-dependent ²H incorporation into *y7* fragment ions (K12K16) and *y12* fragment ions (K8K12K16) from tri-acetylated H4 (4–17) peptides enables quantification of mixed acetylation turnover at H4K12K16 and H4K8K12K16 in tri-acetylated species. **(G)** Estimation of acetylation turnover at H4K12 in di-acetylated H4 (4–17) peptide derived from calculated ²H enrichment differences between positional fragment ions containing these residues.

**Supplementary Figure 5.**
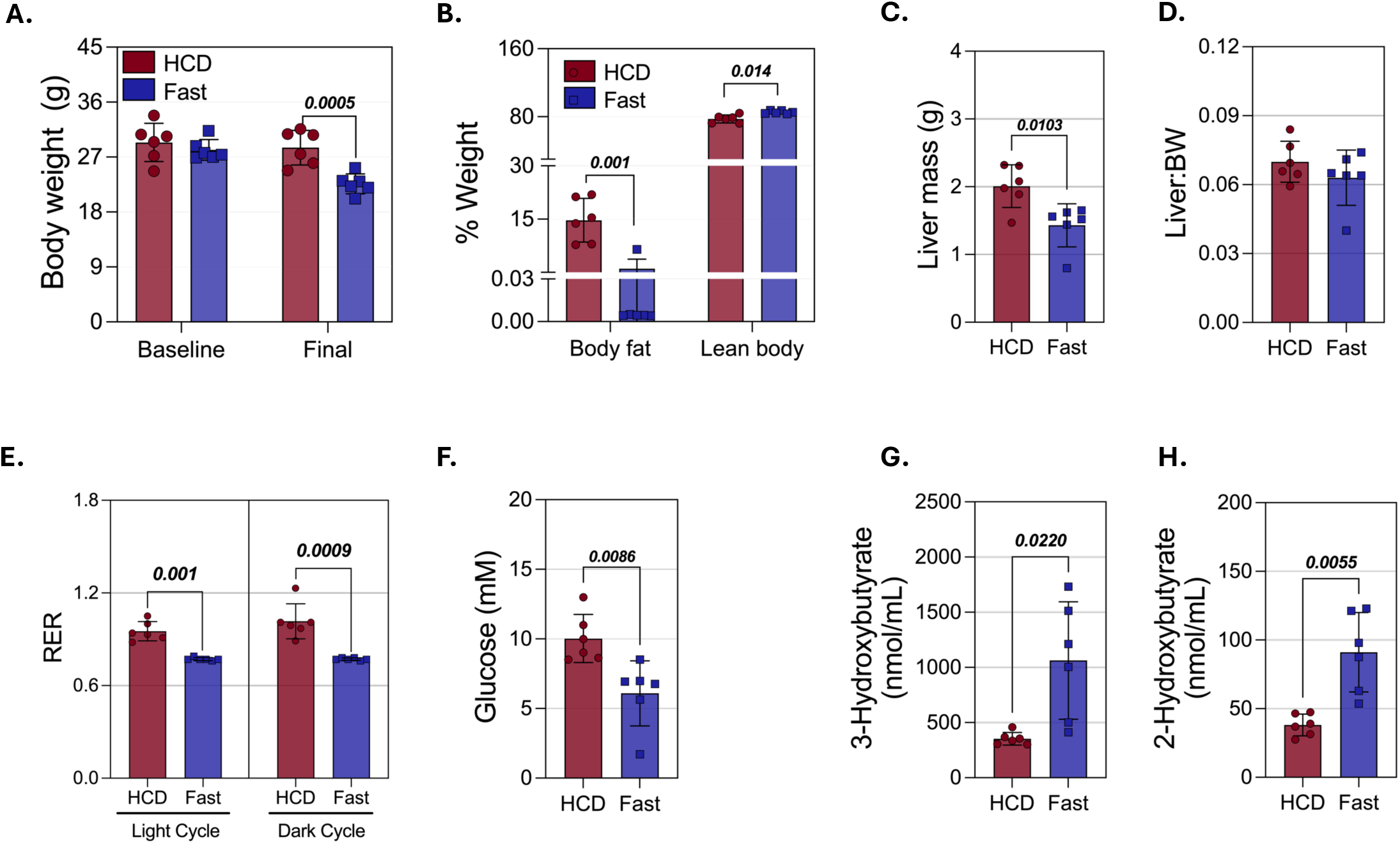
Metabolic characterization of mice under fed (HCD) and fasted conditions. (**A**) Baseline and final body weight, (**B**) body fat mass and lean mass measured by EchoMRI. (**C**) Liver weight and (**D**) liver weight to body weight ratio, measured at the time of tissue collection. (**E**) Respiratory exchange ratio measured by CLAMS metabolic cages. (**F**) Blood glucose, (**G**) plasma 3-hydroxybutyrate, and (H) plasma 2-hydroxybutyrate. Data are presented for mice maintained on a high-carbohydrate diet (HCD) or subjected to fasting. P values were calculated using two-sample t-tests; P < 0.05 was considered statistically significant.

**Supplementary Figure 6.**
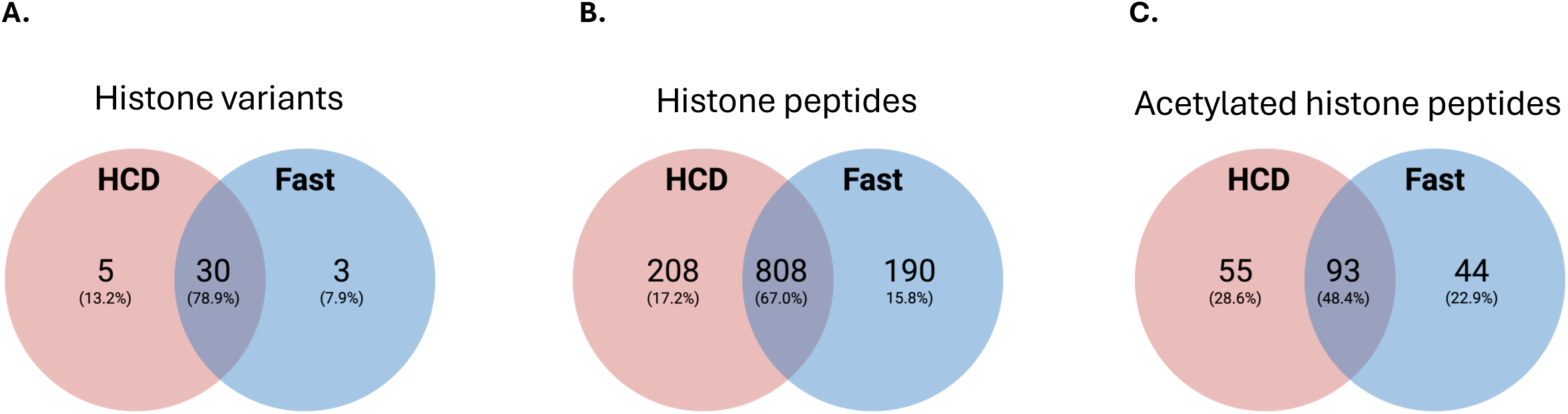
Comparative histone proteomics in fasted and HCD-fed mouse livers. Venn diagrams showing the numbers of shared and unique (**A**) histone variants, (**B**) total peptides, and (**C**) acetylated peptides identified in the livers of fasted and high-carbohydrate diet (HCD)–fed mice.

**Supplementary Figure 7.**
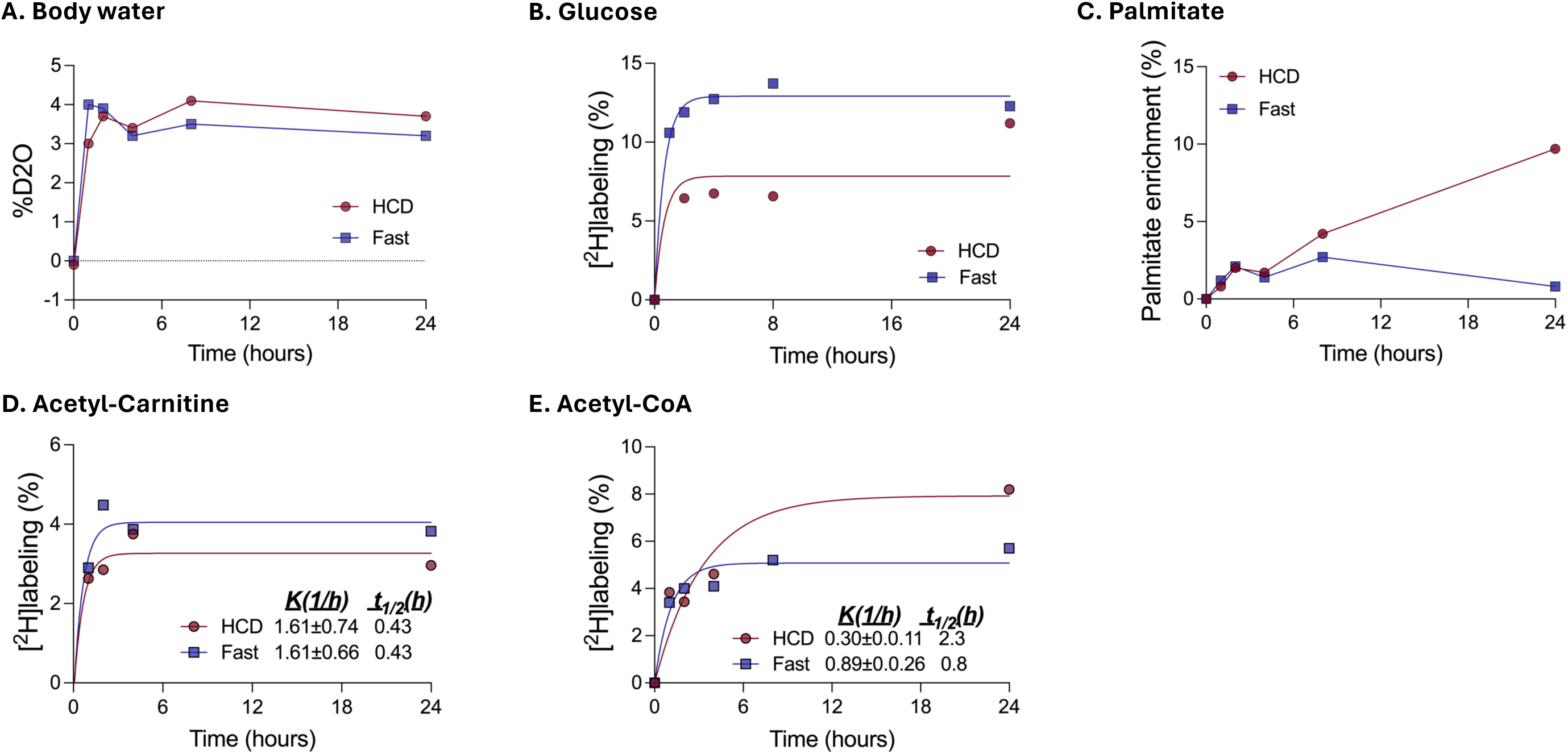
^2^H enrichment of body water and key metabolic intermediates. (**A**) Total body water, (**B**) Glucose, and (**C**) Palmitate. Minimal ²H incorporation into palmitate in fasted mice indicates negligible de novo lipogenesis (DNL) compared with mice fed a high-carbohydrate diet (HCD). (**D**) Acetyl-carnitine, and (**E**) acetyl-CoA. Time-dependent ²H enrichment demonstrates rapid labeling of glycolytic intermediates and acetyl-CoA relative to fatty acids.

**Supplementary Figure 8.**
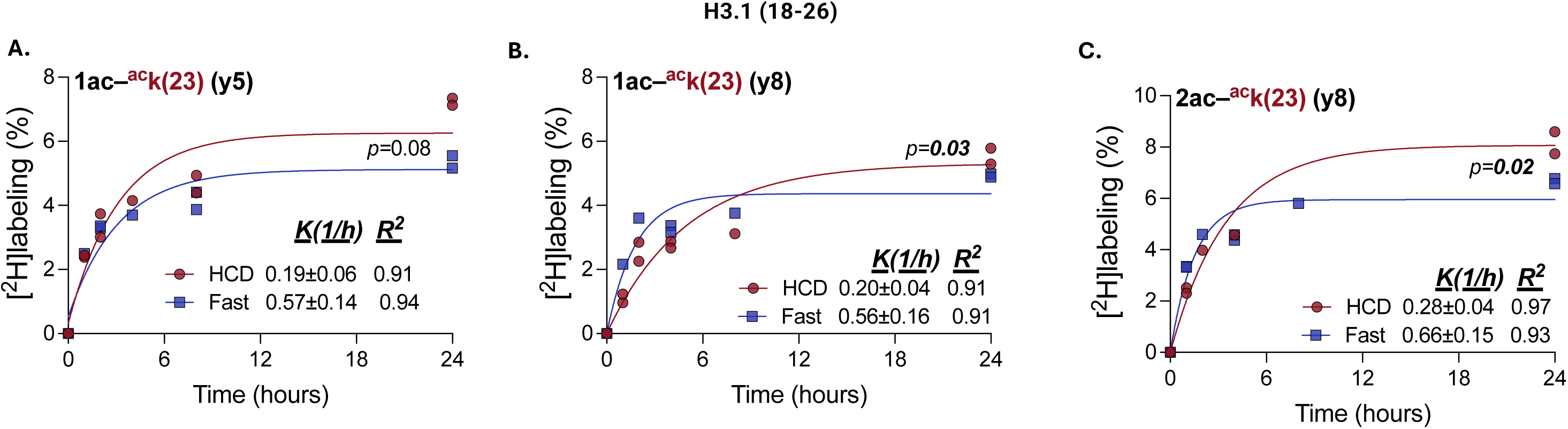
²H labeling of fragment ions from mono- and di-acetylated H3.1 (18–26) peptides in HCD-fed and fasted mouse livers. **(A)** Time-dependent ²H labeling of the *y5* fragment ion from the mono-acetylated **H3.1 (18–26)** peptide containing K23Ac. **(B)** Labeling of the *y8* fragment ion from the mono-acetylated peptide containing K23Ac. **(C)** Labeling of the *y8* fragment ion from the di-acetylated peptide containing K23Ac.

**Supplementary Figure 9.**
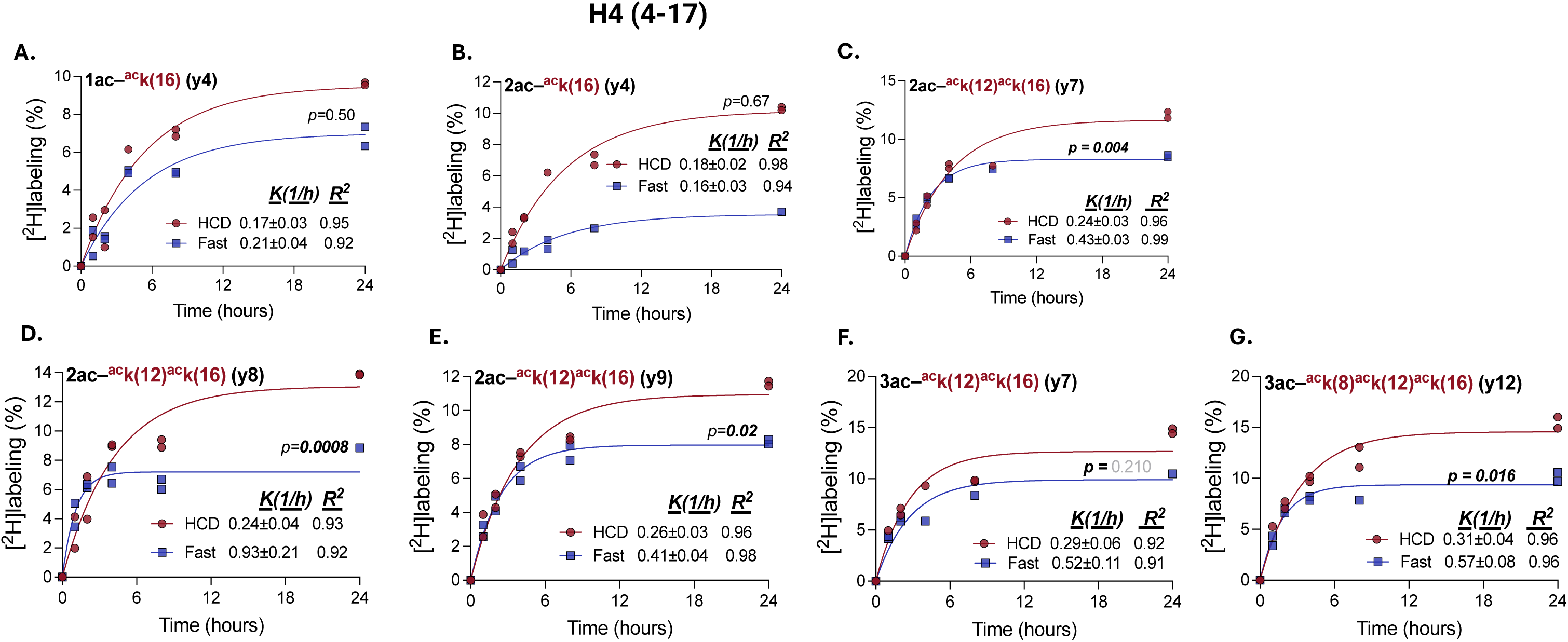
²H labeling of fragment ions from di- and tri-acetylated H4 (4–17) peptides in HCD-fed and fasted mouse livers. **(A)** Labeling of the *y4* fragment ion from the mono-acetylated peptide containing K16Ac. **(B)** Labeling of the *y4* fragment ion from the di-acetylated peptide containing K16Ac. **(C)** Labeling of the *y7* fragment ion from the di-acetylated peptide containing K12Ac and K16Ac. **(D)** Labeling of the *y8* fragment ion from the di-acetylated peptide containing K12Ac and K16Ac. **(E)** Labeling of the *y9* fragment ion from the di-acetylated peptide containing K12Ac and K16Ac. **(F)** Labeling of the *y7* fragment ion from the tri-acetylated peptide containing K12Ac, and K16Ac **(G)** Labeling of the *y12* fragment ion from the tri-acetylated peptide containing K8Ac, K12Ac, and K16Ac. Isotopic enrichment of these fragment ions enables direct estimation of mixed acetylation turnover across multiple lysine residues and was further used to derive indirect site-specific turnover estimates for H4K5 and H4K8 (see main Figure 7).

**Supplementary Figure 10.**
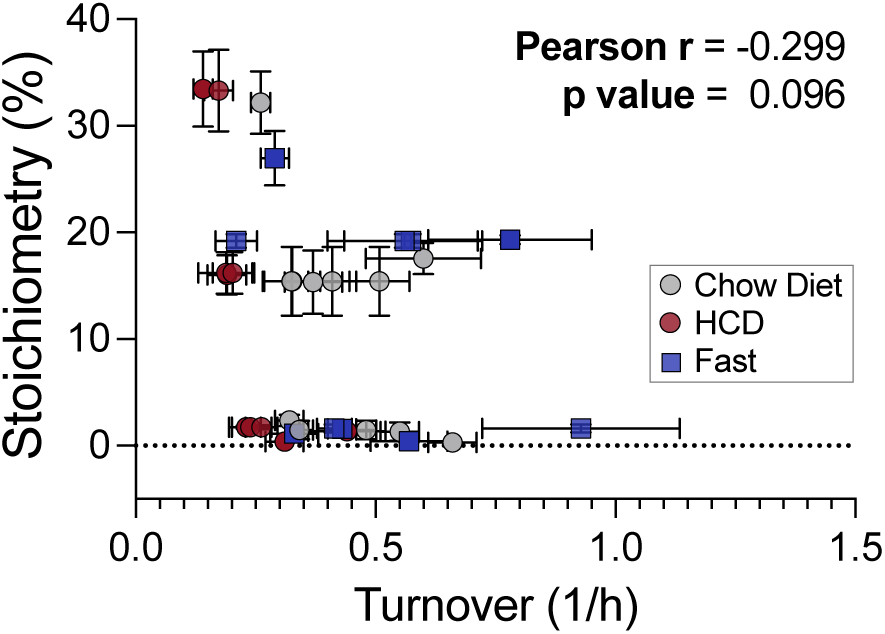
Pearson correlation analysis between site-specific histone acetylation turnover rates and acetylation stoichiometry, illustrating the relationship between dynamic acetylation turnover and steady-state modification levels across quantified histone lysine residues.

## Notes

### Competing Interest Statement

The authors have declared no competing interest.

